# Enhanced bacterial immunity and mammalian genome editing via RNA polymerase-mediated dislodging of Cas9 from double strand DNA breaks

**DOI:** 10.1101/300962

**Authors:** Ryan Clarke, Robert Heler, Matthew S. MacDougall, Nan Cher Yeo, Alejandro Chavez, Maureen Regan, Leslyn Hanakahi, George M. Church, Luciano A. Marraffini, Bradley J. Merrill

## Abstract

The ability to target the Cas9 nuclease to DNA sequences via Watson-Crick base pairing with a single guide RNA (sgRNA) has provided a dynamic tool for genome editing and an essential component of adaptive immune systems in bacteria. After generating a double strand break (DSB), Cas9 remains stably bound to it. Here we show persistent Cas9 binding blocks access to DSB by repair enzymes, reducing genome editing efficiency. Cas9 can be dislodged by translocating RNA polymerases, but only if the polymerase approaches one direction towards the Cas9-DSB complex. By exploiting these RNA polymerase-Cas9 interactions, Cas9 can be conditionally converted into a multi-turnover nuclease, mediating increased mutagenesis frequencies in mammalian cells and enhancing bacterial immunity to bacteriophages. These consequences of a stable Cas9-DSB complex provide insights into the evolution of PAM sequences and a simple method of improving selection of highly active sgRNA for genome editing.

## INTRODUCTION

The clustered regularly interspaced short palindromic repeats (CRISPR) system provides bacteria and archaebacteria an adaptive immune system (Barrangou and Marraffini, 2014). In type II CRISPR systems, immunity begins during the adaptation phase wherein foreign DNA elements near the system’s protospacer adjacent motif (PAM) sequence are recognized, then processed and inserted as the spacers into the CRISPR locus (Barrangou et al., 2007; Garneau et al., 2010; Heler et al., 2015). The immunization phase then begins through expression of the CRISPR loci, and is characterized by spacer transcripts being processed into crRNA (Deltcheva et al., 2011). The crRNA direct Cas9 nuclease activity to foreign DNA by forming a ribonucleoprotein complex with Cas9 and tracrRNA and using the crRNA sequence to identify targets (Jinek et al., 2012). The PAM is an important component that prevents Cas9 from cutting the spacer sequence in its own genome by enabling nuclease activity only when the crRNA target sequence is adjacent to the short DNA sequence also used during the capture of spacers from the foreign DNA (Heler et al., 2015). For repurposing Cas9 to edit gigabase sized genomes, Watson-Crick base pairing of the 5’ 20bp of a single guide RNA (sgRNA) has provided sufficient specificity for widespread use of *S. pyogenes* Cas9 (spCas9) in editing various genomes, including those of mammals (Hsu et al., 2013; Jinek et al., 2013; Mali et al., 2013).

The basic biochemical and biophysical characteristics of spCas9 have been elucidated and exploited for genome editing. The ability to specifically target a single site within the genome without off-target effects has been the focus of considerable research effort (Chen et al., 2017; Kleinstiver et al., 2016; Slaymaker et al., 2016). The relatively minor restrictions the PAM places on genomic sites that can be targeted, and the ease of targeting Cas9 by expressing a short sgRNA have combined to support widespread and pervasive use of Cas9 for genome editing (Barrangou and Doudna, 2016).

In addition to the biochemical properties of Cas9 that provide its target specificity, the nuclease displays other unique properties that distinguish it from non-RNA-guided effector nucleases of bacterial immune systems, such as restriction endonucleases. In contrast to other endonucleases, Cas9 exhibits a remarkably stable enzyme-product state wherein the nuclease remains bound to the double strand break (DSB) it generates (Jinek et al., 2014; Nishimasu et al., 2014; Richardson et al., 2016). The Cas9-DSB state has been shown to persist *in vitro* for ~5.5 hrs (Richardson et al., 2016). Nuclease dead Cas9 (dCas9) and active Cas9 display the same slow off-rate *in vitro* (Richardson et al., 2016). Recent characterization of dCas9 in *E. coli* reported stable binding to target DNA until DNA replication occurs (Jones et al., 2017). Although persistence of Cas9 binding for hour-long periods has not been examined in mammalian cells, fluorescence recovery after photobleaching and single molecule fluorescence studies demonstrated persistence of dCas9 binding during minute-long observations (Knight et al., 2015). Thus the slow off-rate appears to affect Cas9 functionality *in vitro* and *in vivo*.

In contrast to the rapid characterization for how Cas9 targets DNA, the consequences of the persistent enzyme-product state are not understood. The single-turnover characteristic could limit Cas9’s effectiveness when DNA substrates are abundant, such as during phage infection. When DNA substrates are rare, such as when Cas9 is used to edit a unique mammalian genomic sequence, persistence of Cas9-DSB could preclude repair of the DSB by the cell. To date, experimental techniques to manipulate the kinetics of Cas9 dissociation from the DSB have been limited, which has prevented direct analysis of the consequences of the highly stable enzyme-product complex.

In this study, we show that the Cas9-DSB complex can be disrupted by a RNA polymerase transcription activity through the Cas9 target site, but only if the sgRNA of Cas9 is annealed to the DNA strand used as the template by the RNA polymerase. The profound difference caused by the direction of the translocating RNA polymerase enabled examination of the effects of the persistent Cas9-DSB state. Dislodging Cas9 from the DSB stimulates editing efficiency in cells by allowing the ends of the DSB to be accessed by DNA repair machinery. This mechanism causes sgRNA to be more effective if they anneal to the template DNA strand of transcribed genes and also increases the immunity mediated by crRNA that anneal to the template strand of bacteriophage genomes. These data provide insights into the biology of the CRISPR system and provide a simple method of enhancing probability of successful genome editing by choosing sgRNA that anneal to the template strand of DNA.

## RESULTS

### Active transcription through Cas9 target sites increases genome editing frequencies

Several genomic factors that affect genome editing frequencies have been identified with previous studies, including nucleosome occupancy, DNase hyper-sensitive sites (DHSS), and histone marks (H3K4me3) associated with active transcription (Chari et al., 2015; Horlbeck et al., 2016). To complement these findings using a distinct approach, we focused on being able to detect a large range of mutation frequencies as a way to identify genomic variables affecting Cas9-mediated mutagenesis. We examined a collection of 40 sgRNA, each targeting the coding sequence in a different gene (Table S1). Transient transfections were used to express Cas9 and an sgRNA in mouse ES cells, genomic DNA was isolated two days after transfection, and indels were measured by targeted deep sequencing of each genomic target site. Analysis of these 40 sgRNA target sites facilitated robust levels of mutagenesis to be measured, and a wide range of indel frequencies was observed, from 1.5% to 53.7% (Fig. 1A, Table S1). Most sgRNA (33 of 40) displayed a similar and high mutagenesis activity (>30% indel formation). Unexpectedly, the distribution of mutation frequencies was distinctly bimodal, with seven of the 40 displaying substantially lower activity that separated them from the majority of sgRNA (Fig 1A). Interestingly, six of the seven poorly performing sgRNA annealed to the DNA strand that was not used by RNA Pol II as the template for transcription, i.e. the non-template strand (Fig 1B,C, Table S1). The seventh annealed to the template strand of a gene (Actbl2) that was not expressed in ES cells (Fig 1B,C, Table S1). For simplicity, sgRNA that anneal to the non-template strand of a transcribed DNA will hereafter be referred to as “non-template sgRNA”, and those that anneal to the template strand of a transcribed DNA will be referred to as “template sgRNA”.

**Figure 1:**
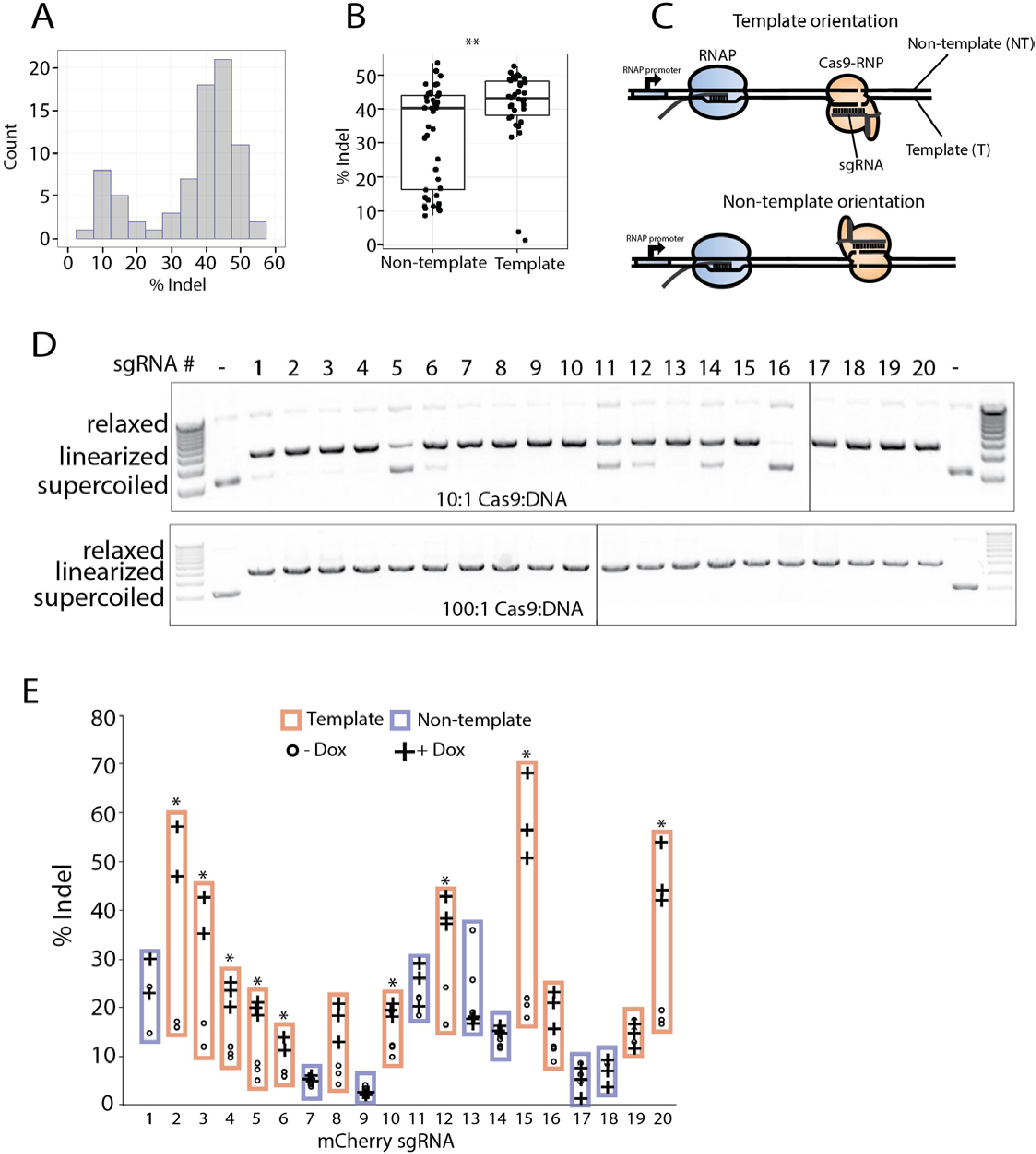
Transcription mediated displacement of Cas9 from the DSB increases genome editing frequencies and is strand dependent. **A,** Bimodal distribution of indel frequencies obtained across 40 distinct mouse genes as measured by targeted deep sequencing 48 hours after transient transfection of Cas9 and sgRNA expression plasmids. Targeted gene, sgRNA sequences, and indel frequencies are presented in Table S1. **B,** Indel frequencies from the **A** were pooled into two groups based on the DNA strand annealing to the sgRNA with respect to the approaching RNA polymerase for each targeted gene, rendering 17 template and 23 non-template sgRNA. Each point represents a mutation frequency of independent transfections, n=2 for each sgRNA. ^∗∗^ = p < 0.01. **C,** Schematic illustrating orientation of Cas9, target DNA, and an approaching RNAP for the two possible RNAP and Cas9 collision orientations: template and non-template. **D,** *In vitro* Cas9 digestions of an mCherry containing plasmid with 20 different sgRNA, tested at two different Cas9:substrate ratios. **E,** Mutagenesis frequencies by sgRNA (#1-20) targeting a doxycycline-inducible mCherry were measured via T7E1 assays (see Fig S2E,F for representative gels). Each point is a biological replicate. ^∗^p<0.05.

To assess the correlation between transcription and indel mutagenesis with additional sgRNA, the large dataset from Chari et al was re-examined. The relationship between indel formation and template/non-template status of sgRNA was tested. Transcription through each targeted site was evaluated by RNA-seq FPKM levels from the same cell line (Fig S1A)(Chavez et al., 2015), and each target gene and its sgRNA were separated into quartiles based on corresponding FPKM values. High levels of gene transcription positively correlates with higher mutation frequency, with a significant ~2 fold difference between quartiles 1 and 4 (Fig S1B, left). This effect from transcription appeared to be caused by increased efficiency from template strand sgRNA (Fig S1B, middle), because transcription levels did not generate statistically significant differences among the bins of non-template sgRNA (Fig S1B, right).

To directly test the effects of transcription through the Cas9 target site, we used a mouse ES cell line harboring a doxycycline (dox) inducible mCherry gene (Fig S1C). Twenty sgRNA (12 template, 8 non-template) targeting the mCherry gene were first assessed for their ability to stimulate Cas9 digestion of DNA in *in vitro* reactions (Fig 1D). All sgRNA were able to stimulate DSB formation in vitro, although five (#5, 11, 12, 14, and 16) required higher concentrations of Cas9 than the other 15 sgRNA (Fig 1D). Mutagenesis frequencies mediated by the 20 sgRNA *in vivo* were measured by T7 endonuclease 1 (T7E1) activities on PCR products generated from genomic DNA isolated two days after transfection of sgRNA and Cas9 expression plasmids (Fig. S1D,E). Without dox-induced transcription of mCherry, the 20 sgRNA displayed a range of indel formation (3%-21%), and the ranges of indel frequencies derived from template sgRNA were not significantly different than non-template sgRNA (Fig. 1E, S1D,E). Stimulating mCherry transcription with dox following transfection did not significantly affect mutagenesis mediated by any of the non-template sgRNA (Fig 1E). By contrast, mutagenesis by 9 of 12 template-strand sgRNA was significantly increased by transcription through the target site (Fig. 1E). The stimulation of mutagenesis caused by transcriptional activity was substantial (2-3 fold) for those sgRNA that were affected. Together, these results show that transcription through a Cas9 target site can stimulate mutagenesis in cells, provided the sgRNA anneals to the DNA strand that serves as the template for the RNA polymerase. We suggest that the transcription-mediated stimulation of mutagenesis precludes template sgRNA from displaying weak indel mutagenesis activity. By contrast, non-template sgRNA are more likely to provide weak activity because they do not benefit from transcription through the target site. Mechanisms underlying this phenomenon are examined below.

### Cas9 precludes double stranded break repair enzymes from accessing DNA ends *in vitro*

Previous biochemical experiments demonstrated that Cas9 remains tightly associated with DNA after generating a DSB (Jinek et al., 2014; Nishimasu et al., 2014; Richardson et al., 2016). Consistent with this property, *in vitro* Cas9 nuclease reactions (as in Fig 1D) required removal of Cas9 with proteinase K in order visualize migration of DNA products into an agarose gel by electrophoresis (Fig 2A). Based on the strength of Cas9 binding to its DSB, we posited that persistence of Cas9 binding to its DSB could be a rate limiting step for mutagenesis in cells. Translocation of RNA polymerase through the Cas9 site could reduce this rate limiting step by destabilizing the enzyme-product complex.

**Figure 2:**
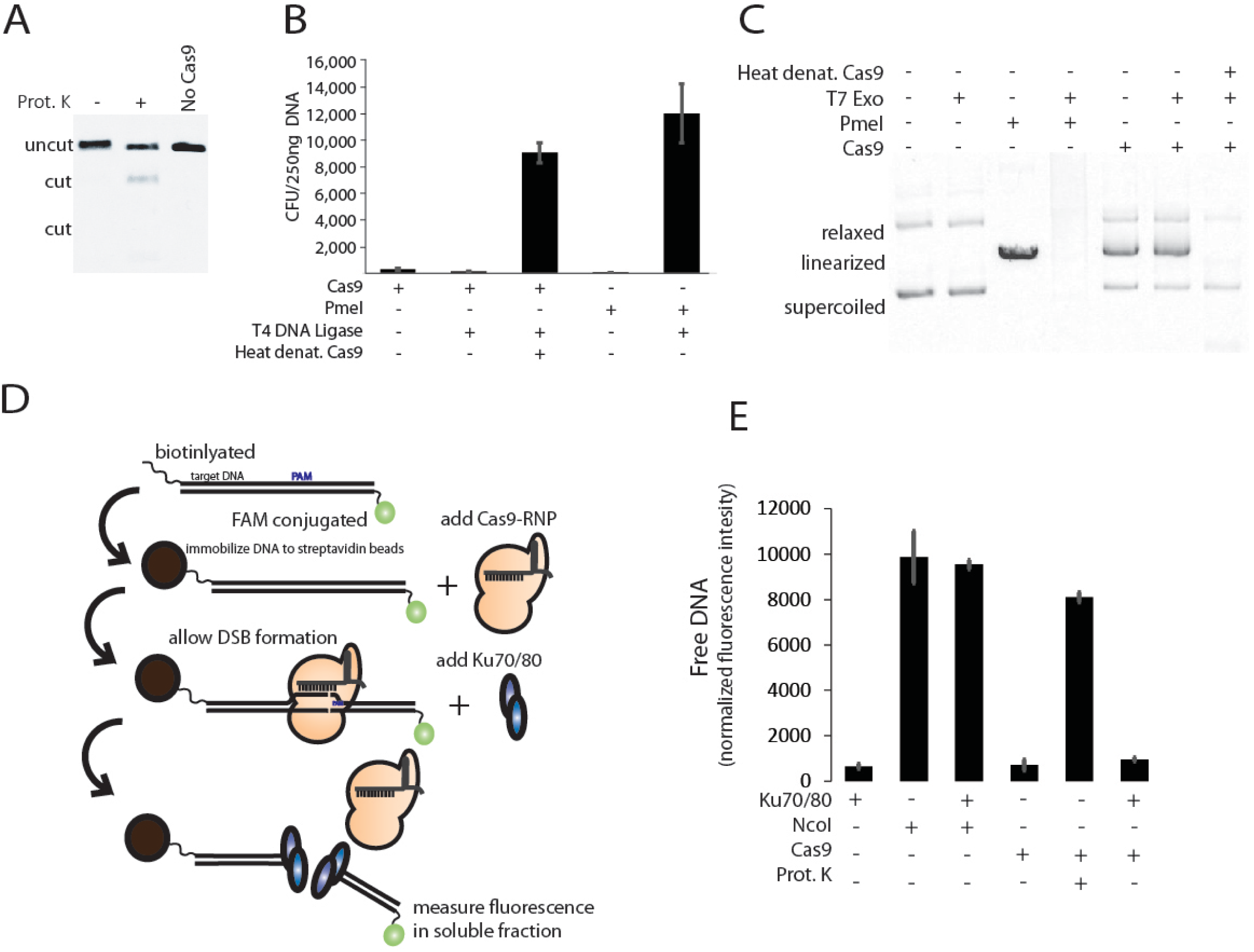
The Cas9-DSB complex precludes DNA repair activities *in vitro.* **A,** Agarose gel electrophoresis of linear dsDNA cut by Cas9, then treated with Proteinase K. **B,** *E. coli* colony formation from circular plasmid DNA undergoing the indicated digestion and ligation conditions. Cas9 was denatured at 75C for 10m before addition of phage T4 DNA ligase. Values represent mean +/- s.d., n=3. **C,** Agarose gel displaying plasmid DNA digested with PmeI restriction endonuclease or Cas9 before incubation with phage T7 exonuclease. **D,** Schematic depicting the experiment in **E** testing if Ku70/80 can displace Cas9 from a DSB. The Cas9-DSB complex is formed on an immobilized, fluorescent substrate then challenged with purified Ku70/80. Disruption of the Cas9-DSB complex causes release of the fluorescent DNA end into the soluble fraction. **E,** Release of fluorescently labeled DNA ends into the soluble fraction after challenging the immobilized target DNA with indicated conditions. NcoI is a restriction endonuclease that cuts the DNA substrate. Values represent mean +/- s.d., n=3.

To begin to test this hypothesis, we determined if persistence of Cas9 binding to DNA prevents DNA end-binding proteins from accessing the Cas9-generated DSB. We tested whether T4 DNA ligase could evict Cas9 from the DSB by first forming Cas9-DSB complexes on a circular plasmid DNA, then adding T4 DNA ligase and incubating at 16°C before using the reactions for bacterial transformation into *E. coli.* A lack of antibiotic-resistant colonies indicated that the ligase was unable to access and repair the Cas9 bound plasmid that encoded ampicillin resistance (Fig 2B). Removing Cas9 by a brief heat denaturation before the ligase reaction restored colony-formation, demonstrating that the Cas9-generated DSB was a competent substrate for T4 DNA ligase if Cas9 was removed from the DNA (Fig 2B). DNA exonuclease activity was examined by comparing degradation of a circular DNA linearized by either a restriction endonuclease or by Cas9 (Fig 2C). Exonuclease activity was prevented at the Cas9-generated DNA ends, unless Cas9 protein was removed by heat denaturation (Fig 2C). These indicate that the persistence of the Cas9-DSB complex prevents the DNA ends from being used as substrates for DNA repair enzymes.

To test whether mammalian DSB end binding proteins could evict Cas9 from its DSB, Cas9 was targeted to a DNA that was immobilized on a bead at one end and fluorescently tagged at the other end. Disruption of the Cas9-DSB complex was detected by measuring soluble fluorescence (Fig. 2D). As a positive control, Cas9 digested DNA was treated with proteinase K to release the fluorescent tag from the bead. When challenging the Cas9-DSB with purified human Ku 70/80, a 100x molar excess of the Ku70/80 complex was incapable of displacing Cas9 from the DSB (Fig. 2E), despite Ku70/80 binding to the other DNA ends present in the reaction (Fig. S2C). Although these *in vitro* observations use a DNA substrate that is not subjected to events occurring on genomic DNA in cells, they demonstrate that the persistent Cas9 binding to DNA can cause the DSB to be inaccessible to DNA end binding proteins. This property is consistent with the possibility that perdurance of Cas9-DSB complex constitutes a rate limiting step during genome editing *in vivo.*

### The Cas9-DSB complex is disrupted by translocating RNA polymerases

We hypothesized that transcription through a Cas9 site increases indel formation, because a translocating RNA polymerase dislodges Cas9 from its DSB (diagramed in Fig 3A). Removing Cas9 from the DSB could stimulate mutagenesis by decreasing the time it takes for the DNA ends to become accessible to cellular repair machinery. To determine if RNA polymerase (RNAP) translocation through the Cas9 site was sufficient to make the DSB accessible to other proteins, we utilized a dsDNA Cas9 substrate harboring the T7 promoter upstream of the cleavage site. The promoter and Cas9 site were orientated so that the sgRNA annealed to the DNA strand that was used as the template by T7 RNAP for transcription. A combined reaction was performed wherein Cas9 digestion of the DNA occurred at the same time as T7 RNAP transcription of the same DNA (Fig 3B). T7 RNAP transcription through the Cas9 site allowed the DSB to be effective substrates and initiate T5 exonuclease activity to degrade the DNA (Fig. 3B). This result indicated that translocation of a T7 RNAP through the Cas9-DSB complex made the DNA ends accessible.

**Figure 3:**
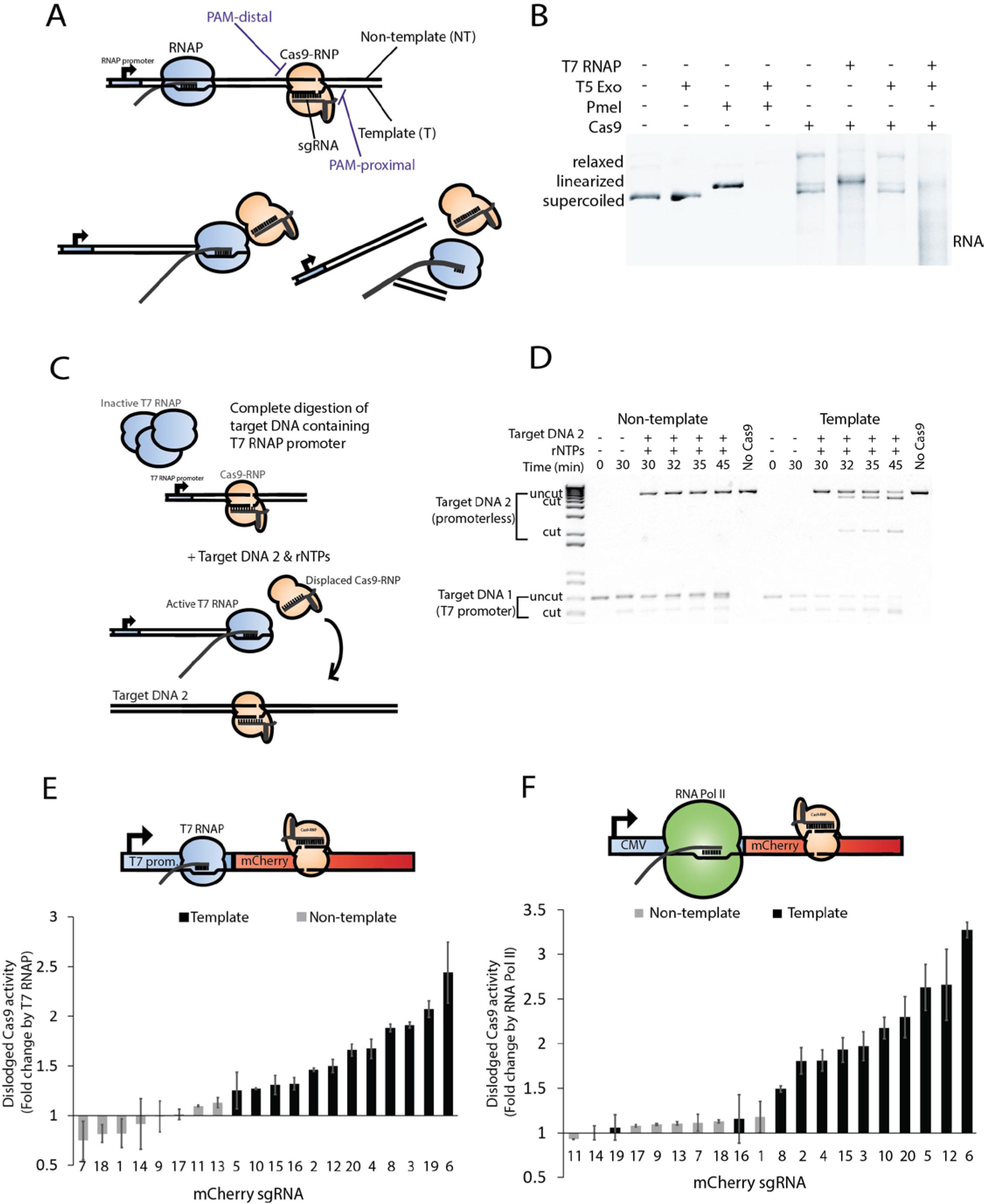
The Cas9-DSB complex is disrupted by translocating RNA polymerases when the sgRNA anneals to the template strand. **A,** Schematic illustrating orientation of Cas9 RNP, target DNA, and T7 RNAP translocation colliding with the PAM-distal surface of Cas9 to disrupt the enzyme-product complex. **B,** Agarose gel displaying plasmid DNA treated with PmeI restriction endonuclease or Cas9 before incubation with T5 exonuclease and/or T7 RNA polymerase (T7 RNAP). Prot. K: proteinase K. **C,** Schematic illustrating experiment to detect T7 RNAP displaced Cas9 molecules. Target DNA 2 lacks T7 promoter sequence. Target DNA 1 and target DNA 2 each have the same DNA sequence targeted by Cas9. **D,** Agarose gel displaying Cas9 cleavage of target DNA with T7 RNAP promoter on either end of the DNA to generate template or non-template collisions for the same sgRNA sequence. The promoterless target DNA 2 serves as the substrate for Cas9-RNPs that were displaced by T7 RNAP as illustrated in **C**. **E,** Fold change Cas9 activity dislodged from mCherry DNA by T7 RNAP activity as measured by the soluble fraction fluorescent levels for a fluorescent, promoter-less target DNA. T7 RNAP activity was controlled by addition of rNTPs. See experiment schematic in Fig. S2D. **F,** Fold change Cas9 activity dislodged from mCherry DNA by RNA Pol II activity from nuclear extracts. Activity was measured by the soluble fraction fluorescent levels for a fluorescent, promoter-less target DNA. RNA Pol II activity was controlled by addition of α-amanitin. See experiment schematic in Fig. S2D. Error bars represent mean +/- s.d., from n=2.

The DNA strands emanating from one side of the Cas9-DSB complex display more freedom than DNA from the opposite side. Notably, DNA at the PAM-distal surface of Cas9 (Fig 3A) have been shown to be accessible to hybridize with complementary ssDNA; whereas DNA at the PAM-proximal surface of Cas9 does not hybridize with complementary ssDNA probes (Richardson et al., 2016). The 5’ to 3’ direction of RNA polymerization causes a translocating RNAP to collide with PAM-distal surface of the Cas9-DSB when the sgRNA anneals to the DNA strand used as a template by RNAP (as displayed in Fig 3A). Conversely, when the sgRNA anneals to the non-template strand, translocation of the RNAP will result in a collision with the PAM-proximal surface of the Cas9-DSB complex. We hypothesized that these differences could affect genome editing *in vivo* if the orientation of the collision affected the ability of RNAP to disrupt the Cas9-DSB complex.

To test a strand bias in the ability of RNAP to dislodge Cas9, we developed an assay that took advantage of the dislodged Cas9-RNP possibly being able to bind to another DNA molecule and generate a DSB in that DNA as long as it contained the sgRNA target sequence. First, Cas9 and a T7 promoter-containing target DNA (target DNA 1) were incubated (30 min) to allow DNA cleavage and formation of the Cas9-DSB complex. Next, a promoterless target DNA (target DNA 2) containing an identical Cas9 target site was added (Fig 3C). Note that a ten-fold molar excess of target DNA 1 relative to Cas9 and stability of the Cas9-DSB complex combined to prevent detectable cleavage of target DNA 2 in the absence of transcription (Fig. 3D). Transcription through the Cas9-DSB complex in target DNA 1 was activated by adding rNTPs, and we began measuring cleavage of target DNA 2 after two minutes of transcription. Target DNA 2 was cut rapidly after initiating transcription, but only if the the sgRNA bound to target DNA 1 was annealed to the template strand (Fig 3D, right side). Collision with Cas9-DSB in the non-template orientation did not generate nuclease activity on target DNA 2 (Fig 3D, left side). Since target DNA 2 was not transcribed in this assay, the stimulation of its digestion by T7 RNAP could not be caused by a differential activity of Cas9 on actively transcribed DNA, per se. The rapid digestion of target DNA 2 after RNAP activation is most consistent with RNAP activity on target DNA 1 removing Cas9 from its DSB, and allowing it to digest another DNA molecule. Finally, the inability of T7 RNAP to stimulate target DNA 2 digestion in the non-template sgRNA orientation is consistent with Cas9-DSB complexes being resistant to dissolution by RNAP colliding with the PAM-proximal surface of Cas9. Together these data indicate that the Cas9-DSB complex can be disrupted by RNAP, if the sgRNA anneals to the template strand.

We examined whether the strand biased ability to displace Cas9 *in vitro* was a general phenomenon by measuring displacement levels for the 20 sgRNA targeted across a linear mCherry substrate. Reactions were performed in the presence or absence of rNTPs to compare transcription mediated displacement levels for each sgRNA. Displacement of Cas9 activity from a T7-conatining mCherry DNA was measured using an immobilized, fluorescently tagged target DNA 2. After completion of the combined digestion/transcription reaction, displacement was assessed by fold change in soluble fluorescence stimulated by RNAP (Fig. 3E, S2D). These reactions showed that all template annealed sgRNA were compatible with displacement by T7 RNAP (Fig 3E). In contrast, all of the non-template sgRNA were recalcitrant to displacement (Fig 3E).

T7 RNAP and mammalian RNA Pol II can be considered very different from each other in terms of their biophysical and biochemical properties. Since the *in vitro* results elucidated above used T7 RNAP, but we propose that the *in vivo* genome editing effects of transcription are cause by RNA Pol II, the ability of RNA Pol II to displace Cas9 from its DSB were determined. A fluorescent displacement assay was performed essentially as described above for the T7 RNAP experiment (Fig 3E, S2D); however, target DNA 2 was used to detect Cas9 dislodged off of a CMV-mCherry template by RNA Pol II activity from mouse ES cell nuclear extracts (Fig. 3F, S2E). To determine dependence of transcription for Cas9 displacement, reactions were performed in the presence or absence of the RNA Pol II/III inhibitor, α-amanatin, (Fig. S2E). Fold changes in fluorescence levels revealed that none of the eight non-template sgRNA were significantly displaced (Fig 3F). Thus, the non-template sgRNA prevented displacement of Cas9 from DSB for either RNAP tested. By contrast, 10 out of 12 template sgRNA were substantially displaced by RNA Pol II activity (Fig 3F). Interestingly, template sgRNA displayed varying levels of displacement in both transcription scenarios, suggesting sgRNA-determined variability in disruption of the Cas9-DSB complex. Notably, two template sgRNA (#16,19) were not displaced by RNA Pol II activity, and a third (#8) displayed a low level of displacement relative to other template sgRNA. Levels of indel mutagenesis with these three template sgRNA did not significantly increase after transcriptional activation of mCherry *in vivo* (Fig 2E). Together, these data indicate that a strand biased RNA Pol II displacement of Cas9 from its DSB stimulates indel mutagenesis in cells.

### RNA polymerase can convert Cas9 into a multi-turnover nuclease

When using Cas9 for genome editing in cells or organisms, the nuclease is typically expressed or delivered at a high molar ratio relative to its DNA substrates, which are often only 2-4 copies per cell. As such, efficiency of genome editing is likely less dependent on the capabilities of one Cas9 nuclease to processively digest many DNA substrates than it is on a rapid detection of the DSB by the cell’s repair machinery. However, when RNAP collides with the Cas9-DSB complex, the displaced Cas9 molecule retained its nuclease activity (Fig 3B,D), suggesting that Cas9 could be converted from a single-turnover nuclease to a multi-turnover nuclease. An ability of a single Cas9 molecule to digest many DNA substrates could be important when saturating levels of DNA targets need to be digested, such as when high multiplicities of infection occur during bacteriophage infection.

To determine the multi-turnover capabilities of Cas9, a twofold excess of a single, T7 promoter-containing target DNA was used as a substrate for *in vitro* Cas9 digestion reactions. To test template and non-template orientations using the same sgRNA, the promoter was placed on either end of the target DNA. After an initial 30 min digestion of half of the DNA, addition of rNTPs was used to initiate T7 RNAP activity, and RNAP-stimulated cleavage of DNA was measured for up to 30 min (Fig. 4A). Placing the T7 promoter so that the sgRNA annealed to the template strand stimulated Cas9 cleavage activity with rapid kinetics similar to those observed at the start of a reaction (Fig. 4A, S3A). No stimulation was observed with the non-template strand orientation (Fig. 4A). Continual displacement of Cas9 by T7 RNAP did not appear to disrupt the Cas9-sgRNA interaction, because Cas9 did not exchange sgRNA molecules after being displaced (Fig. S3B).

**Figure 4:**
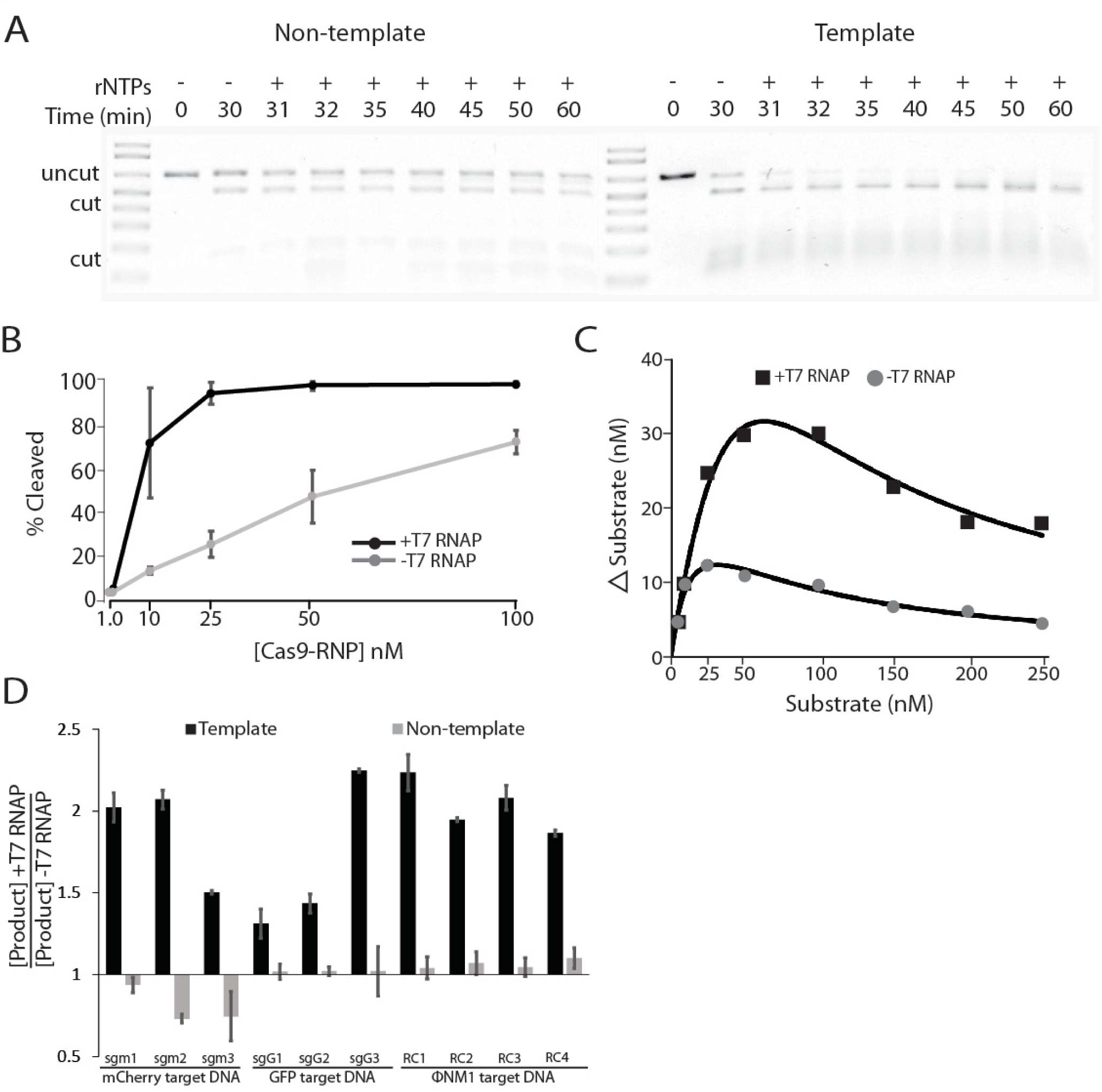
Strand dependent ability of translocating T7 RNA polymerase to stimulate *in vitro* multiturnover nuclease activity by Cas9. **A,** Multi-turnover nuclease activity occurs exclusively in the template strand orientation. Agarose gel displaying hybrid Cas9 digestion and transcription reaction. T7 promoter was placed on either end of the target DNA to achieve template or non-template orientation. Transcription was initiated at 30min by addition of rNTPs. **B,** Titration of Cas9 in the presence or absence of T7 RNAP. Target DNA was held constant at 150nM. Values represent mean +/- s.d., n=3. (representative gel shown in Fig. S3C). **C,** Titration of substrate in the presence or absence of T7 RNAP. Cas9 was held constant at 12.5nM. Values represent mean +/- s.d., n=2. (representative gel shown in Fig S3D). **D,** Cas9 digestion reactions of target DNAs harboring T7 RNAP promoters in the template or non-template strand orientation. See Fig. S4A for schematics and Fig S4B,C and Fig S6 for representative gels. Values represent mean +/- s.d. of fold change in cleaved DNA caused by addition of T7 RNAP, n=3.

Altering the amount of Cas9 (Fig. 4B) or the amount of T7-promoter-containing DNA substrate (Fig. 4C) in a reaction revealed substantial capabilities of Cas9 to function as a multi-turnover nuclease *in vitro.* Diluting Cas9 showed that T7 RNAP increased the capacity for template sgRNA orientation by 10-fold compared to reactions without T7 RNAP translocation through Cas9, which functioned as a single-turnover nuclease (Fig. 4B, S3C). The improved capacity increased kinetics of Cas9 activity at saturating substrate concentrations (Fig. 4C). T7 RNAP converted Cas9 to a multi-turnover nuclease for a variety of sgRNA and target DNAs tested, but only when the sgRNA annealed to the template DNA strand (Fig. 4D, S4A-C, S6). The magnitude of stimulation by T7 RNAP varied among template strand sgRNA, but it did not appear to correlate with GC content of the target site (Fig. S4D) or the GC content in the sequence next to the PAM (Fig. S4E). In summary, when combined with T7 RNAP and sgRNA in the template strand orientation, Cas9 was effectively transformed from a single-turnover enzyme into a multi-turnover enzyme.

### PAM sequences and protospacer targets are more frequently located on template strand of *Streptococci* phages

We wondered whether the strand bias in Cas9’s potential to act as a multi-turnover nuclease contributed to bacterial immunity. Given a stoichiometry of multiple bacteriophage particles infecting individual bacterial cells, we reasoned that Cas9 functioning as a multi-turnover nuclease could have substantial benefits over a single-turnover nuclease. A multi-turnover nuclease could significantly enhance bacteriophage immunity by allowing a single Cas9 molecule to destroy more than one bacteriophage genome. Therefore, we examined whether there were differences in the frequencies of Cas9 predicted to act as a single-turnover versus multi-turnover nuclease on bacteriophage genomes.

Interestingly, the nucleotide composition of bacteriophage genomes differ in the DNA strand replicated by leading strand versus lagging strand DNA synthesis (Jin et al., 2014; Kwan et al., 2005; Lobry, 1996; Uchiyama et al., 2008). This phenomenon has been named GC skew, and underlying causes for it remain uncertain. For Streptococcus phages that infect *S. pyogenes* and *S. thermophilus*, the GC skew is reflected in the nucleotide composition of the plus strand (34% adenine : 27% threonine, and 22% guanine : 17% cytosine). The structure of these bacteriophage genomes places the transcription of genes in predominantly one direction; thus, template strands have a different nucleotide composition than non-template strands. Consequently, the potential PAM sites for spCas9 (NGG) and *S. thermophilus* Cas9 (stCas9 -NNAGAAW) are not strand neutral. Instead, they preferentially target the template strand at about a 2:1 ratio for spCas9 and 3:1 ratio for stCas9 (Fig. 5A, B, S5A). Mapping crRNA identified from bacteriophage insensitive mutant strains to bacteriophage genomes showed that the actual frequency of crRNA annealing to the template strand are more abundant than those annealing to the non-template strand (Fig. S5B, Table S3) (Achigar et al., 2017; Levin et al., 2013). Rational engineering of Cas9 proteins showed that mutagenesis can relatively simply change the PAM sequence Cas9 recognizes (Kleinstiver et al., 2016), indicating that the nuclease has potential to be preferentially targeted to either or neither strand in bacteriophage genomes. Based on the above correlations, we hypothesized that targeting Cas9 to anneal to the bacteriophage template strand provides a selective advantage by allowing Cas9 to function as a multi-turnover nuclease during active transcription through target sites.

**Figure 5:**
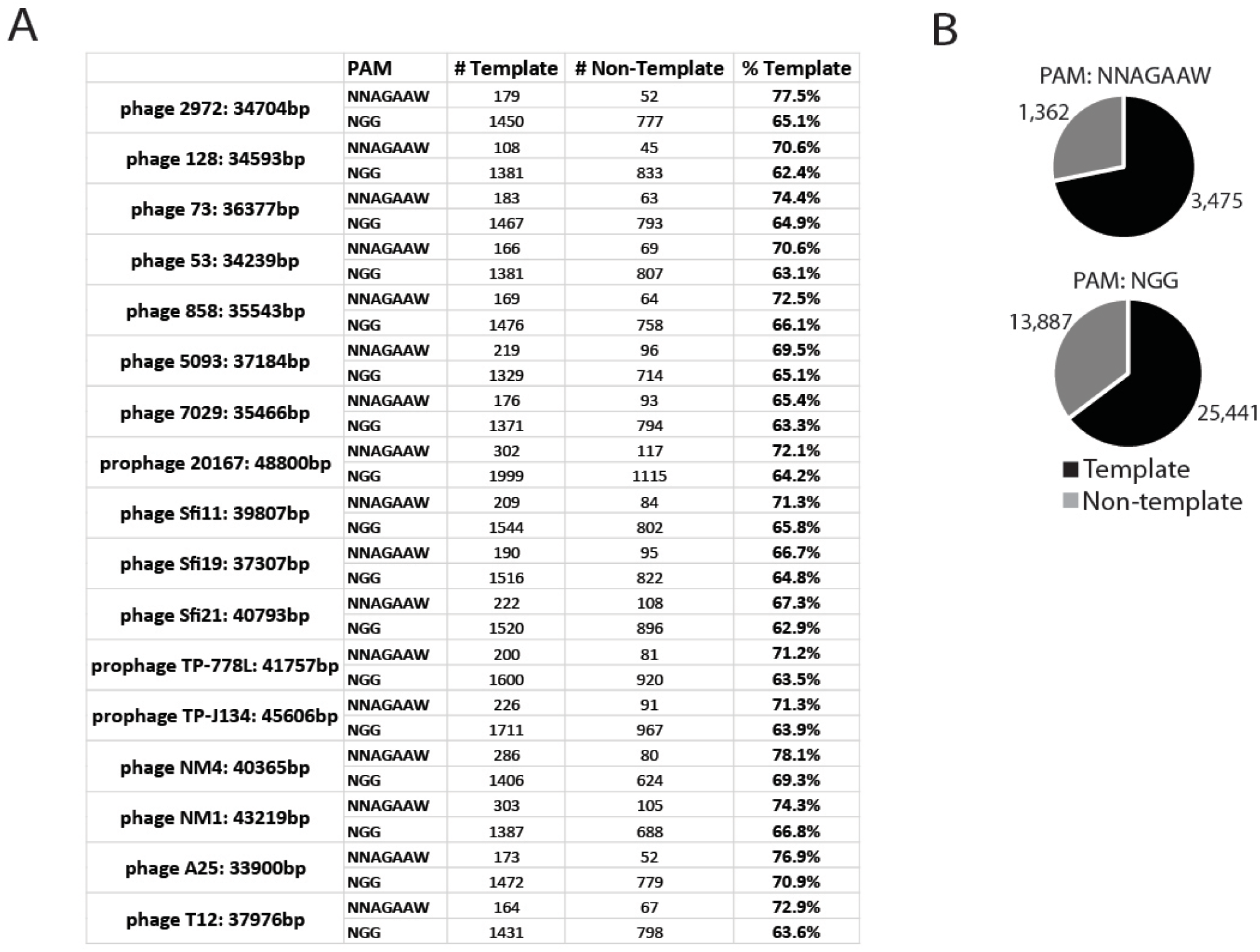
PAM sequences across *Streptococci* phage are more frequently oriented on the template strand. **A,** Of the 16 surveyed *streptococcus* phages, all harbor majority of PAM sequences on DNA strand corresponding to the template strands for RNA transcription. NNAGGAW = *S. thermophilus* PAM and NGG = *S. pyogenes* PAM. **B,** Distribution of all PAM sequences among genomes analyzed in **A**.

### Template targeted protospacers enhance bacterial adaptive immunity

To directly test a strand bias effect on bacterial immunity, we used two virulent versions of the ΦNM1 phage. One contains a mutation that inactivates the promoter required for transcription of the lysogeny cassette (ΦNM1γ6) (Goldberg et al., 2014). The other expresses the lysogeny cassette, but it harbors an inactivating deletion within the cl repressor gene (ΦNM1h1) (Fig. 6A). Therefore, neither phage can establish lysogeny, but they differ in the transcription of the lysogeny cassette.

**Figure 6:**
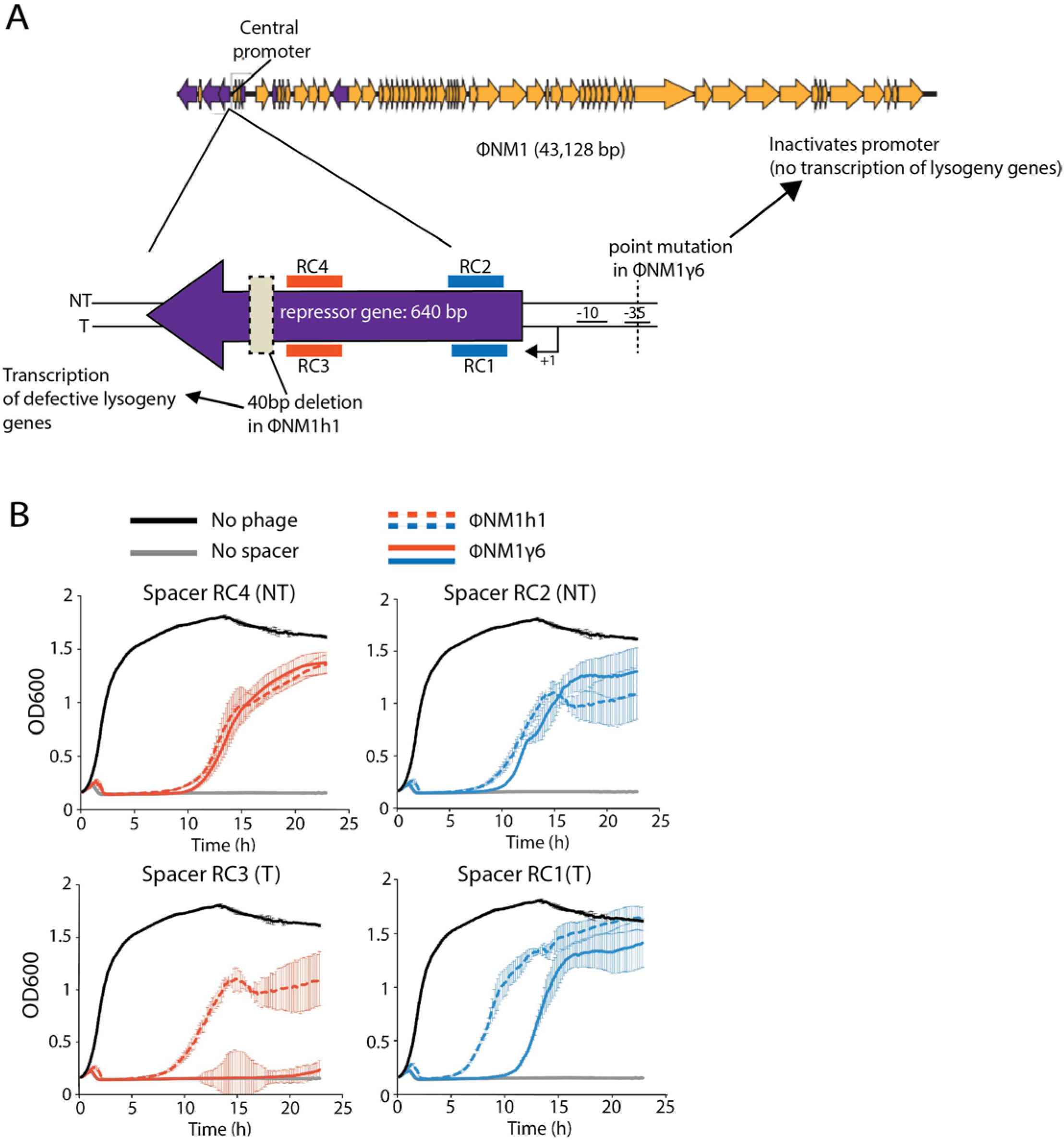
Template strand-targeted protospacers enhance phage interference. **A,** General organization of the phage ΦNM1 genome and targeting strategy of the lysogenic repressor gene in the ΦNM1h1 and ΦNM1γ6 mutant phages. RC1 – RC4 indicate sites for crRNA used in panel **B**. **B,** Growth curves of *S. aureus* strains harboring spacers RC1-4 after infection with ΦNM1h1 or ΦNM1γ6. T: template strand orientation. NT: non-template strand orientation.

To test the effect of transcription through a Cas9 target site, we generated different bacterial strains harboring spacers annealing to either template or non-template strand sequences within the repressor gene found in both ΦNM1γ6 and ΦNM1h1 (Fig. 6A). Each strain was infected with each phage, and their survival was determined by measuring OD600 over time (Fig. 6B). The interference efficiency of each spacer against the two phages was interpreted from plate-reader growth curves of infected bacterial cultures. The two spacers targeting the non-template strand (RC2 and RC4) showed similar interference against either phage regardless of whether transcription was active (ΦNM1h1) or inactive (ΦNM1γ6). On the contrary, spacers targeting the template strand (RC1 or RC3) were notably more effective at providing immunity against the actively transcribed target (ΦNM1h1) than the inactive target (ΦNM1γ6). The same four target sites within ΦNM1 were tested for the ability of T7 RNAP translocation to turn Cas9 into a multi-turnover nuclease *in vitro* (Fig. S6), demonstrating the template strand bias effect on the phage genome. These results show that active transcription across Cas9 targets improves CRISPR immunity by converting Cas9 into a multi-turnover enzyme, but the effect appears to be restricted to Cas9 annealed to the template strand.

## DISCUSSION

The consequences of the persistent Cas9-DSB state were elucidated by identifying conditions that dissociate Cas9 from its DNA products. The stable enzyme-product complex precludes DNA repair activities *in vitro*, but it can be disrupted by translocating RNA polymerases in a strand biased manner, conditionally converting Cas9 into a multi-turnover nuclease. This dislodging from the DSB had significant effects on genome editing and bacterial immunity, by increasing mutation frequencies in mammalian cells and mediating enhanced phage interference through multi-turnover nuclease activity.

This new understanding of the interaction between Cas9 and RNA polymerases can be directly applied to CRISPR-Cas9 based genome editing procedures. Our sample of 20 sgRNA to the same gene demonstrates that all these sgRNA are competent to mediate Cas9 digestion of substrates *in vitro*, yet they displayed substantial variability for indel frequency *in vivo.* Genomic factors, such as nucleosome occupancy, have previously been shown to affect indel frequency (Horlbeck et al., 2016); however, they are unlikely to affect variability here because all sgRNA targeted a single locus, which should not vary in any of the previously identified factors. Instead a large degree of variability among sgRNA was clearly attributable to the direction of RNA pol II translocation through the Cas9 target site. Although the 2-3 fold increased mutagenesis should be considered substantial, more benefit to genome editing will likely be gained by reducing the probability of using a so-called “dud” sgRNA by avoiding non-template sgRNA. Subsequent research resulting in the modification of Cas9 or discovery of small molecules that destabilize the Cas9-DSB complex could stimulate CRISPR-Cas9 based mutagenesis, especially at non-transcribed sites and in cells with low DNA metabolic activity. In the absence of such advances, our findings provide a simple and straight forward path for increasing efficiency of Cas9-mediated mutagenesis, which is to preferentially use only sgRNA that anneal to the template strand.

This strand biased removal of Cas9 from its DSB is interesting to consider alongside recent biochemical analyses of dCas9 dissociation from DNA. The DNA emerging from the PAM-proximal surface of Cas9 is double stranded and is not accessible to exogenous ssDNA for strand invasion (Richardson et al., 2016). Because of the RNA:DNA hybrid between the sgRNA and target DNA, the DNA emerging from the PAM-distal surface is single stranded, and ssDNA hybridization to PAM-distal sequence can displace dCas9 from its target (Jinek et al., 2014; Nishimasu et al., 2014; Richardson et al., 2016). Mismatched base pairing had the greatest effect on dCas9 dissociation when located at PAM-distal positions, suggesting that the 5’ end of the guide RNA contributes most significantly to the Cas9 off-rate (Boyle et al., 2017). RNA polymerases approaching the PAM-distal surface of the Cas9-DSB complex should have freedom to collide with Cas9. We suggest that a physical collision from RNA polymerases dislodges Cas9 from the DSB, facilitating repair of the DSB, and enabling the Cas9 molecule to cut an additional target DNA. The GC content of the target site and sequence adjacent to the PAM did not significantly affect displacement of Cas9 from the DSB for either orientation. Further biophysical studies are needed to determine why some template sgRNA were more affected by RNA Pol II translocation than others.

Multiple studies have shown that after the CRISPR-Cas9 immune response, some of the acquired viral spacers are highly represented in the population of surviving bacteria (Heler et al., 2015; Paez-Espino et al., 2013). Most likely multiple factors determine the success of a new spacer, but it is tempting to speculate that one such factor could be the disposition of the target sequence with respect to its transcription. Our results suggest that spacers leading to the engagement of Cas9 with its target in a disposition where the nuclease can be removed by RNAP after cleavage would allow a more efficient cleavage of the often multiple phage genomes infecting the host. Such spacers would mediate a more robust immune response, and therefore would be positively selected from the pool of all the acquired spacers. It is possible that the strand biased PAM sequences of StCas9 and SpCas9 evolved to target the strand of bacteriophage genomes where it can become multi-turnover. In comparing evolution of PAM sequences and bacteriophage genomes, it should be noted that the distribution of PAM sequences on either strand of the bacteriophage genome may differ among bacteriophage that infect a given bacteria. However, in the organisms examined here, *Steptococci* and their associated bacteriophage, the PAM sequence used by Cas9 to target the more effective strand is relatively simple, and altering it requires only a small number of mutations (Kleinstiver et al., 2015). By contrast, the GC-skew is pervasive over the entirety of the bacteriophage genome and is effectively unchangeable relative to the PAM sequence. We propose that targeting the bacteriophage template strand provides an advantage because it will more frequently result in multi-turnover nucleases upon transcription of lytic genes.

## AUTHOR CONTRIBUTIONS

R.C. and B.J.M. jointly designed the study; R.H. and L.A.M. conceived the phage experiments. R.C., R.H., M.S.M. and M.R. designed and performed experiments; L.H., G.M.C., and M.R. provided reagents; R.C., and M.S.M. conducted analysis of mutation frequency with technical advice and support from N.C.Y., and A.C. R.C., R.H., L.A.M., and B.J.M. wrote the manuscript with significant advice and discussion from all authors.

## ACKNOWLEDGMENTS

We would like to thank Ying Su and Dr. Arnon Lavie for assistance with the purification of Cas9, Dr. Sylvain Moineau for assistance with bacteriophage genomics, Dr. Stefan Green and the DNA Services core facility at UIC for assistance with sequencing, Dr. Miljan Simonovic for his assistance with interpretation of *in vitro* experiments. This work was supported by the National Institutes of Health [R01-HD081534 to B.J.M; 1DP2AI104556 to L.A.M.], and the UIC Center for Clinical and Translational Sciences [R.C., M.S.M.]. R.H. is the recipient of a Howard Hughes International Student Research Fellowship. L.A.M. is supported by the Rita Allen Scholars Program, a Burroughs Wellcome Fund PATH award, and a HHMI-Simons Faculty Scholar Award, N.Y.C. and G.M.C. are supported by a US National Institutes of Health National Human Genome Research Institute grant no. RM1 HG008525 and the Wyss Institute for Biologically Inspired Engineering, A.C. is supported by a Burroughs Wellcome Fund CAMS award.

## STAR METHODS

### Contact for Reagent and Resource Sharing

Further information and requests for reagents can be directed to, and will be fulfilled, by the corresponding author Bradley J. Merrill.

### Experimental Model and Subject Details

#### Cell culture

Mouse embryonic stem (ES) cells harboring the Rex1:EGFPd2 insertion (A gift from Dr. Austin Smith) (*24*), or a Rosa26::TetOn-Otx2-mCherry insertion(A gift from Dr. Shen-his Yang) (*25*) were maintained 10cm dishes previously coated with 0.1% gelatin in Knockout DMEM media supplemented with the following: 15% KnockOut Serum Replacement, 2mM l-Glutamin, 1mM HEPES, 1× MEM NEAA, 55μM 2-mercaptoethanol, 100 U/ml LIF, and 3μM CHIR99021. Cell cultures were routinely split 1:10 with 0.25% trypsin–EDTA every 2–3 days.

### Method Details

#### Recombinant Cas9 purification

Cas9 (pMJ806, Addgene #39312) was expressed and purified by a combination of affinity, ion exchange and size exclusion chromatographic steps as previously described (*20*).

#### sgRNA synthesis for *in vitro* Cas9-RNP

All sgRNAs were cloned into pSPgRNA (Addgene, #47108) following the protocol optimized for pX330 base plasmids (https://www.addgene.org/crispr/zhang/) (*21*). sgRNA oligo sequences, listed without BbsI sticky ends used for cloning, can be found in Supplementary Figures S1H, S2A, and S5D. Templates for *in vitro* transcription were generated via PCR mediated fusion of the T7 RNAP promoter to the 5’ end of the sgRNA sequence using the appropriate pSPgRNA as the reaction template DNA. PCR reactions were performed using Phusion high GC buffer (NEB) and standard PCR conditions (98°C for 30s, 30 cycles of 98°C for 5s, 64°C for 10s and 72°C for 15s, and one cycle of 72°C for 5m). PCR products were then column purified (Qiagen) and eluted in TE (10mM Tris-HCl pH 8.0, 1mM EDTA). DNA concentrations were determined using a Nanodrop 2000 (ThermoFisher Scientific), and then were diluted to 200nM when used as templates for *in vitro* transcription reactions. The transcription reactions contained 5.0μg/ml purified recombinant T7 RNAP (a gift from Dr. Miljan Simonovic) and 1x transcription buffer (40mM Tris-HCl pH8.0, 2mM spermidine, 10mM MgCl2, 5mM DTT, 2.5mM rNTPs). Following incubation at 37°C for 1 hour, reactions were treated with RNase free DNase I (ThermoFisher Scientific) and column purified using the Zymo RNA Clean & Concentrator kit following the manufacturer’s protocol. The purified RNA products were eluted from the column in 15μl of water.

#### DNA templates for *in vitro* Cas9 nuclease reactions

Linear target DNAs for hybrid digestion and transcription assays: mouse Lef1, mCherry, and GFP target DNAs were generated by PCR amplification using 50ng genomic DNA from mESC Rosa26::TetON-Otx2-mCherry cells in a reaction using Phusion high GC buffer (NEB) and standard PCR conditions (98°C for 30s, 30 cycles of 98°C for 5s, 64°C for 10s and 72°C for 15s, and one cycle of 72 °C for 5m). ΦNM1 genomic DNA was amplified with the same parameters, except using Phusion HF buffer. All PCR products were column purified (Qiagen), eluted in TE, and concentrations were determined with a Nanodrop 2000 (ThermoFisher Scientific). For experiments testing effects of T7 RNAP on Cas9, the DNA template was a segment of the mouse Lef1 gene generated with primer set #1 (Table S4), unless otherwise stated.

Plasmid target DNAs: For reactions that required circular dsDNA templates (experiments testing accessibility of exonuclease or ligase enzymes), plasmid target DNAs were prepared using TOPO TA cloning. PCR products of the previously described Lef1::PGK-neo and Ctnnb1::EGFP DNA sequences (*22*) were cloned into the pCR4-TOPO Vector (ThermoFisher Scientific).

#### *In vitro* Cas9 DSB formation assays

The basic Cas9 DSB formation assay was prepared in 1x Cas9 digestion buffer (40mM Tris, pH8.0, 10mM MgCl2, 5mM DTT) with a final concentration of 100nM Cas9, unless otherwise stated. Prior to addition of DNA templates, sgRNA was added in molar excess, and incubated at room temperature for 10 min to ensure formation of the Cas9-RNP. Target DNA was added to a final concentration of 200nM and a final reaction volume of 50μl, unless otherwise stated. Reactions were incubated at 37°C for 25 min, then either heat inactivated at 75°C for 10 min or treated with Proteinase K at 37°C for 15 min. DNA fragments from portion of each reaction (usually 15μl) were separated by electrophoresis 1.5% agarose gel, and visualized with ethidium bromide staining.

For reactions involving T7 RNAP transcription, basic Cas9 digestion conditions were applied, except 1x transcription buffer was used, unless otherwise stated. Upon addition of the target DNA, T7 RNAP was added to a final concentration of 5.0μg/ml. Reactions were placed at 37°C for 25 min, unless otherwise stated, and then heat inactivated at 75°C for 10 min. DNase free RNase A (NEB) was added to all reactions except Fig S4A and S5E, then incubated at 37°C for 30 min before separating DNA fragments on a 1.5% agarose gel.

T7 and T5 exonuclease assays were performed in 1x Cas9 digestion buffer, unless otherwise stated. T7 exonuclease assays was performed with the Lef1::PGK-Neo plasmid and digested using sgLef1. T5 exonuclease assays were performed with Ctnnb1::EGFP plasmid and digested using sgG2. T5 exonuclease assays containing T7 RNAP were performed in 1x transcription buffer. All reactions contained 100nM Cas9:RNP, 200nM target DNA, and 10U of the appropriate exonuclease. Reactions were subject to Proteinase K treatment before loading onto a 1% agarose gel.

T4 DNA ligase and Cas9 digestion assays were performed in T4 DNA Ligase buffer containing ATP (NEB). Cas9 containing reactions were performed with 200nM Cas9:RNP (sgLef1) and 100nM Lef1::PGK-Neo plasmid were allowed to incubate for 30 min at 37°C, then the temperature was lowered to 16°C and 40U of T4 DNA ligase (NEB) was added and allowed 30 min of incubation. Reactions were transformed into competent DH5a in 3 serial dilutions, and ampicillin-resistant colony forming units determined following overnight incubation at 37°C.

Titrations of Cas9 or substrate: Cas9 hybrid digestion and transcription reactions were performed using sgG2 and a GFP target DNA generated with primer set #6 (Table S4). Cleavage frequencies were measure using ImageJ.

#### Ku70/80 competition assay

Recombinant human Ku70/80 was purified as previously described (*23*). A 5’ biotinylated primer (5’ *BIOSG*-GCCTCACACGGAATCT 3’) and a 3’ FAM conjugated primer (5’ GAGAGCCCTCTCCCAATCTTC-FAM 3’) (Integrated DNA Technologies) were used to amplify a 650bp Lef1 target DNA, PCR products were column purified (Qiagen), and eluted in TE. MyOne Dynabeads (ThermoFisher) were prepared as described by the manufacturer to immobilize 750ng of target DNA to ~4μl of beads. Cas9 and sgRNA were pre-incubated in 1x Cas9 digestion buffer (40mM Tris, pH8.0, 10mM MgCl2, 5mM DTT) for 30 min at room temperature, added to the immobilized DNA in a 5:1 molar ratio, and incubated for 25 min at 37°C. Control reactions without Cas9, but containing DNase, NcoI, and/or Ku70/80 were prepared simultaneously and incubated for 25 min at 37°C. Reactions containing Cas9 were then subject to Proteinase K treatment or addition of excess Ku70/80 (100 fold excess), and incubated for 15 min at 37°C. Bead-bound DNA fragments were then collected by placing reaction tubes on a magnet, and 10μl of the soluble fraction was transferred to a 384 well plate in technical triplicates. FAM fluorescence levels were measured using a Tecan Infinite Pro200. Calculations were made after subtracting the background fluorescence levels of reactions containing the immobilized but uncleaved FAM labeled DNA. Three independently set up reactions were performed for each reaction condition.

#### Mammalian nuclear extract preparation

mESC were grown in a 10cm dish to 10×10^6^ confluency and scraped into 3 ml PBS, then pelleted at 16,000 rpm for 10 min at 4°C. The supernatant was then aspirated and the pellet was resuspended in 800 ul of ice cold Buffer A (10mM HEPES pH 7.9, 1.5mM MgCl2, 10mM KCL, 0.5mM DTT, and 1% protease inhibitors). The pellet was incubated on ice for 10 min, vortexed for 10 sec, then centrifuges at 4°C at 4,000 rpm for 10 min. The supernatant (cytoplasmic fraction) was discarded, and the pellet was resuspended in 200μl of Buffer B (10mM HEPES pH 7.9, 0.4mM NaCl, 10mM KCL, 1.5mM MgCl2, 0.1mM EDTA, 12.5% glycerol, 0.5mM DTT, and 1% protease inhibitors). The resuspended pellet was incubated on ice for 30 min then centrifuged at 14,000 rpm for 20 min at 4°C. The supernatant (nuclear fraction) was aliquoted and stored at -80°C.

#### Nuclear extract transcriptional activity validation (qPCR)

Nuclear extract (6μl per reaction) was added to reactions containing: 12μl 1x NE transcription buffer (20mM HEPES pH 7.9, 100mM KCL, 0.2mM EDTA, 0.5mM DTT, 20% glycerol), 3μl of 50mM MgCl2, 1.2μl 25mM rNTPs, and 38nM CMV-mCherry (serving as RNA Pol II transcription template, see below for PCR amplification procedure) to create final reaction volumes of 45μl. Control reactions contained 6μg of α-amanitin (Sigma-Aldrich #04622) or lacked rNTPs. Reactions were incubated at 37°C for 45 min, then 10U of DNase I was added (ThermoFisher) and incubated at 37°C for 30 min. After completion of DNA digestion, 150μl of TE and 200μl of 25:24:1 phenol:chloroform:isoamyl (Sigma-Aldrich) were added. Reactions were then vortexed for 15 sec, briefly centrifuged, then the aqueous layer was transferred to a fresh 1.5 ml tube. RNA was precipitated from the mixture by adding 3 volumes of ice cold 100% ethanol, then centrifuged for 10min at top speed. RNAs were reconstituted in water and diluted to 500ng/μl. 1μg of RNA was converted to cDNA using Superscript III (ThermoFisher) and quantitative real time PCR was performed using primer set #10 (Table S4) by combining 250ng of cDNA from each sample was with Perfecta SYBR Green Supermix (Quanta #95053). qPCR was performed on a C1000 thermal cycler and CFX96 Real Time System (Bio-Rad) with the following parameters: 95°C for 2 min, then 40 cycles of 95°C for 30s and 60°C for 45s. Quantities were normalized to control reactions of CMV-mCherry DNA used to create a standard curve. Standard log transformation of Ct values and standard curve equation was then applied before calculating fold change of experimental conditions over the no rNTP condition.

#### Fluorescent Cas9-RNP displacement assay

Generation of RNA Pol II or T7 RNAP promoter containing target DNAs: CMV-mCherry was amplified with primer set #11 (Table S4) to include the polyA from pmCherry-C1 (Clontech), and T7-mCherry was amplified with primer set #4 (Table S4). PCRs conditions contained Phusion high HF buffer (NEB) and standard PCR conditions (98°C for 30s, 30 cycles of 98°C for 5s, 64°C for 10s and 72°C for 20s, and one cycle of 72°C for 5m), and PCR products were column purified (Qiagen), and eluted in TE.

Generation of displaced Cas9 fluorescent detection substrates: Fluorescent target DNAs (FT-DNA) were generated to contain 3 or 4 Cas9 target sites per FT-DNA to accommodate all 20 mCherry sgRNA, rendering DNAs that range from 134bp to 90bp (Table S2). FT-DNAs were prepared by ordering single stranded sense ultramers (IDT) and PCR amplifying with a 5’ biotinylated primer (5’ *BIOSG*-CGTAAACGGCCACAAGTTCAG 3’) and a 3’ FAM conjugated primer (5’ CTTGTACAGCTCGTCCATGCC-*FAM* 3’). PCR conditions consisted of Phusion high HF buffer (NEB) and standard PCR conditions (98°C for 30s, 30 cycles of 98°C for 5s, 61°C for 10s and 72°C for 5s, and one cycle of 72°C for 5m), and PCR products were column purified (Qiagen), and eluted in TE.

Displacement assays with mammalian nuclear extracts: To test the effect of RNA polymerase II, Cas9 digestion reactions were carried out in 15μl reactions containing: in 4μl of 1x NE transcription buffer, 1μl of 50mM MgCl2, 0.4μl 25mM rNTPs, and 2.1μl of freshly thawed nuclear extract. Cas9 was added to a final concentration of 26nM and respective sgRNA was added in excess, then incubated at RT for 10 min to allow formation of RNP. After formation of the RNP, 6μg of α-amanitin was added to respective reactions, then CMV-mCherry was added to a final concentration to all reactions to a final concentration of 38nM and to render a final volume of 15ul for all reactions. Reactions were then incubated at 37°C for 45 min. While the hybrid transcription/digestion reactions were incubating, FT-DNAs were immobilized to MyOne Dynabeads (ThermoFisher) as described by the manufacturer. The immobilized FT-DNAs were heated at 75°C for 5 min to remove non-specific binding, then washed twice, then resuspended in 1x NE transcription so FT-DNA was at a concentration of 100ng/μl. Upon completion of the Cas9 transcription/digestion reactions, the bead:FT-DNA conjugates were added to each reaction so FT-DNAs were in 2:1 molar ratio to CMV-mCherry. The reactions were incubated at 37°C for 15min, then heated at 75°C for 10 min to denature the displaced Cas9 which was bound to FT-DNAs thereby releasing the cleaved fluorescent end of the FT-DNAs into the soluble fraction. All reactions were then placed on a magnet, and the soluble fraction was removed and placed into a suitable plate for reading FAM fluorescence levels were measured using a Tecan Infinite Pro200. Calculations were made after subtracting the background fluorescence levels of reactions containing the immobilized but uncleaved FT-DNAs respectively. Two independently set up reactions were performed for each reaction condition.

Displacement assays with T7 RNA polymerase: Reactions were performed in the exact manner as the mammalian nuclear extract displacement assays except with minimal changes: reaction buffer was 1x transcription buffer, and presence or absence of transcription was controlled through presence or absence of rNTPs rather than using α-amanitin. Two independently set up reactions were performed for each reaction condition.

#### Transfection and selection conditions

Within 2 hrs of transfections, 0.25 × 10^5^ ES cells were freshly plated in each well of 24 wells dishes. For each well, 2.5μl of Lipofectamine 2000 and relevant DNAs were incubated in 125μl OPTI-MEM (GIBCO #31985) before adding to wells. For the Cas9 mutagenesis of 40 distinct genes in ES cells, transfections included 150ng pPGKpuro (Addgene plasmid # 11349), 150ng pX330 (lacking sgRNA insert), and 150ng of the relevant pSPgRNA plasmid. To assess background mutation rate due to possible deep sequencing or amplification errors, a transfection containing pSPgRNA with empty sgRNA site was assessed alongside the other sgRNA-containing transfections. Two days after transfection, cells were split into 2μg/ml puromycin and selection was applied for 48hrs before isolating genomic DNA by overnight lysis with Bradley Lysis buffer (10mM Tris-HCl, 10mM EDTA, 0.5% SDS, 10mM NaCl) containing 1mg/ml Proteinase K, followed with EtOH/NaCl precipitation, two 70% EtOH washes, and eluted in 50μl of TE. For mCherry targeting, transfections contained the same DNA, except pSPgRNA targeted the mCherry genomic insertion, genomic DNA was isolated 48 hours after transfection in 50μl of Quick Extract solution (Epicentre) for T7E1 assays.

#### T7 endonuclease 1 assays

Genomic DNA was used as a template in a PCR reaction using Phusion polymerase (NEB) and standard PCR conditions (98°C for 30s, 30 cycles of 98°C for 5s, 55°C for 10s and 72°C for 25s, and one cycle of 72°C for 5m). 5μl of each PCR product was added to 19μl of 1x NEBuffer 2 (NEB), denatured at 95C for 10 min, then brought down to room temperature by decreasing the temperature 1C per second. 1μl of T7E1 (NEB) was added to each reaction, and allowed to incubate at 37°C for 25 min. DNA fragments were separated by electrophoresis through a 1.5% agarose gel. Gel images were analyzed and indel frequencies were quantified using ImageJ.

#### Flow cytometry

Single-cell suspensions were prepared by trypsinization and re-suspension in 2% FBS/PBS/2mM EDTA. Cells were analysed on a LSRFortessa flow cytometer. Data analysis was performed using FlowJo v9.3.2. Live cells were gated by forward scatter and side scatter area. Singlets were gated by side scatter area and side scatter width. At least 5 × 10^5^ singlet, live cells were counted for each sample. mCherry fluorescence events were quantified by gating the appropriate channel using fluorescence negative cells as control.

#### Targeted deep-sequencing preparation

Preparation:genomic DNA was harvested four days after transfection and approximately 100ng of DNA was used in PCR to amplify respective target sites while attaching adapter sequences for subsequent barcoding steps (Table S1 for NGS primers). PCR products were analyzed via agarose gel and then distinct amplicons were pooled for each replicate respectively in equal amounts based on ImageJ quantification. Pooled PCR products were purified with AMPure beads (Agilent), and 5ng of the purified pools was barcoded with Fluidigm Access Array barcodes using AccuPrimer II (ThermoFisher Scientific) PCR mix (95°C for 5m, 8 cycles of 95°C for 30s, 60°C for 30s and 72°C for 30s, and one cycle of 72°C for 7m). Barcoded PCR products were analyzed on a 2200 TapeStation (Agilent) before and after 2 rounds of 0.6x SPRI bead purification to exclude primer dimers. A final pool of amplicons was created and loaded onto an Illumina MiniSeq generating 150bp paired-end reads.

#### Generation of spacers targeting ΦNM1

Plasmids harboring Cas9, tracrRNA and single-spacer arrays targeting ΦNM1 were constructed via BsaI cloning onto pDB114 as described previously (*17*). Specifically, spacers RC1 (plasmid pRH320), RC2 (pRH322), RC3 (pRH324) and RC4 (pRH326) were constructed by annealing oligo pairs H560-H561, H564-H565, H568-H569 and H572-H573, respectively. Each pair of annealed oligos contains compatible BsaI overhangs and can be found in Table S3.

#### ΦNM1 infection assays

Phage ΦNM1h1 was isolated as an escaper of CRISPR type III targeting of ΦNM1 with spacer 4B (*16*). Plate reader growth curves of bacteria infected with phage were conducted as described previously (*16*) with minor modifications. Overnight cultures were diluted 1:100 into 2ml of fresh BHI supplemented with appropriate antibiotics and 5mM CaCl2 and grown to an OD600 of ~0.2. Immune cells carrying targeting spacers were diluted with cells lacking CRISPR-Cas in a 1:10,000 ratio and infected with either ΦNM1h1 or ΦNM1g6 at MOI 1. To produce plate reader growth curves, 200μl of infected cultures, normalized for OD600, were transferred to a 96-well plate in triplicate. OD600 measurements were collected every 10 min for 24 hrs.

### Quantifications

#### Agarose gel quantifications

For all Cas9 digestion reactions and T7E1 assays, percent cleavage values were determined by measuring densitometry of individual DNA bands in ImageJ, then dividing the total cleaved DNA by total DNA.

#### Targeted deep-sequencing analysis

Determination of indel frequencies made use of CRISPResso command line tools that demultiplexed by amplicon, where appropriate, and then determined indel frequency by alignment to reference amplicon files (*28*). Outputs were assembled and analyzed using custom command-line, python, and R scripts which are available upon request.

#### Bioinformatic analysis of RNA seq vs indel frequencies

The source of large scale indel mutagenesis and RNA-seq data were from previously published reports (*9*, *26*). Blat and bedtools command line tools (*27*) were used to classify each of the sgRNA used by Chari et al (*9*) as targeting either the template or non-template gene strand. All data were merged and visualized using RStudio version 1.0.136 (package: ggplot2), allowing for the determination of the effect of FPKM and strand orientation on indel frequency.

### Statistical Analysis

#### Agarose gel quantifications for T7E1

The three biological replicates of the mutagenesis data presented in Fig. 1E were analyzed for statistical significance using RStudio. Statistical analyses were performed by generating p values for each sgRNA with a two sample t-test to compare plus and minus doxycycline, then all p values were adjusted via Bonferroni correction.

#### Targeted deep-sequencing comparisons

Statistical analyses were performed by pooling indel frequencies for all sgRNA annealing to the template or non-template strand, creating two separate groups. Then, unpaired, two tailed t-test to was performed.

#### Bioinformatic analysis of RNA seq vs indel frequencies

Statistical analyses and significance were determined with Multiple Comparisons of Means with Tukey contrasts (package: multcomp).

### Data and Software Availability

All scripts for statistical analysis or preparation of NGS data mentioned throughout the methods section are available upon request.

Raw sequencing data will be available on Mendeley upon time of publication.

## SUPPLEMENTARY FIGURES AND LEGENDS

**Figure S1.**
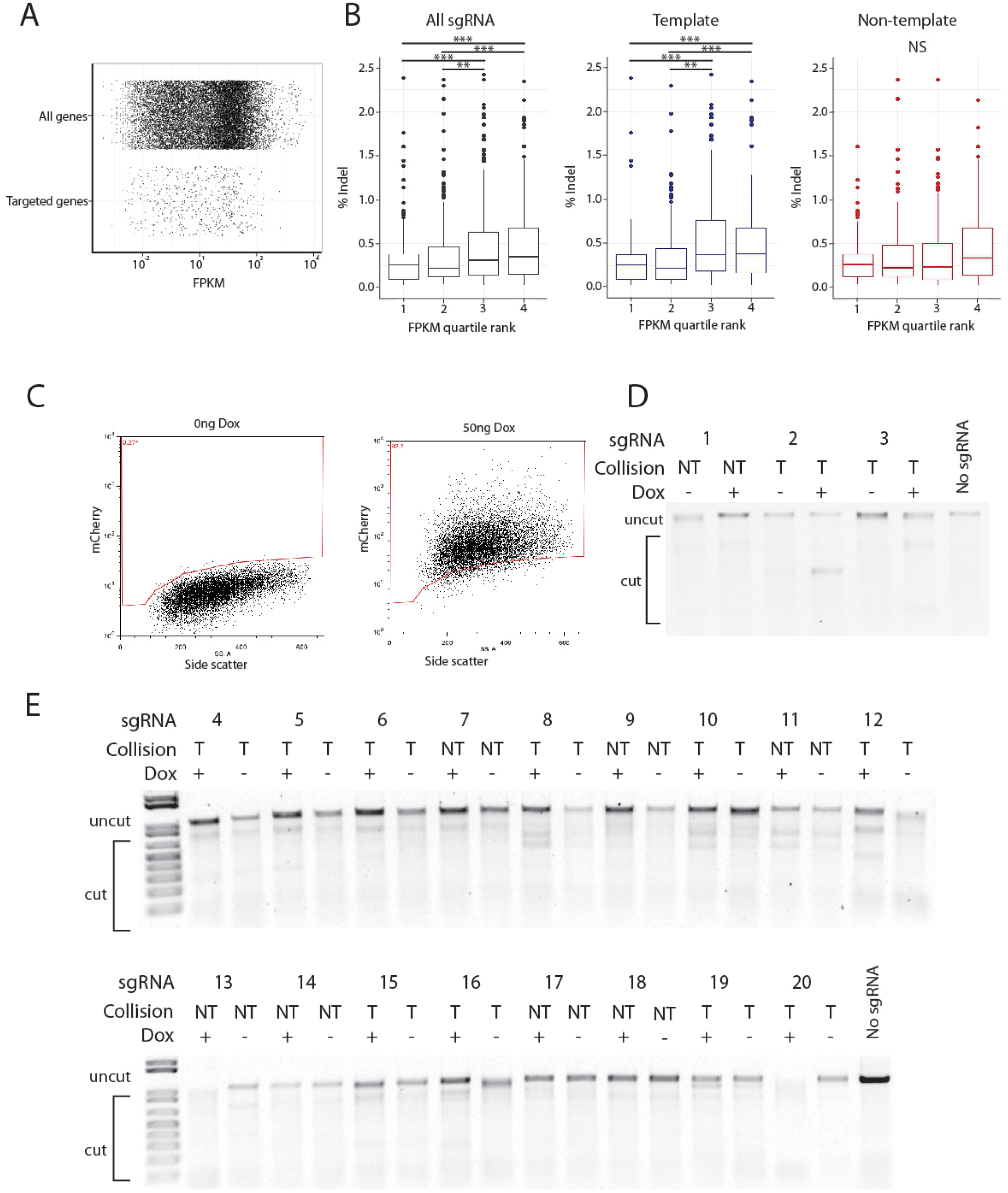
Related to Figure 1: Effects of strand bias on *in vivo* mutagenesis frequency. **A**, Distribution of RNA seq FPKM values for the 975 targeted genes compared to the entire HEK293 genome. Distribution of all detected transcripts by gene in HEK293T cells (20,096 genes) and corresponding sgRNA-targeted genes (975) previously reported in Chari et al, 2015. **B**, RNA levels positively correlate with indel frequencies among 975 sgRNA targeting the human genome. FPKM quartile rank: bins of sgRNA were assembled by their associated RNA-seq values (1-4 = low to high). “All” sgRNA plot contains all sgRNA used in study (243 to 244 sgRNA per bin). “Template” plot contains sgRNA annealing to template strand only (120 to 121 sgRNA per bin. “Non-template” plot contains sgRNA annealing non-template strand only (122 to 123 sgRNA per bin). ^∗∗^ = p < 0.01, ^∗∗∗^ = p < 0.001. **C**, Representative flow cytometry analysis displaying induction of mCherry fluorescence after treating with 50ng/ml doxycycline (dox) for 48 hours. **D,E**, Representative agarose gels displaying T7E1 reactions presented in Fig 1E. (E) n = 2 and (F) n = 3 replicates.

**Figure S2.**
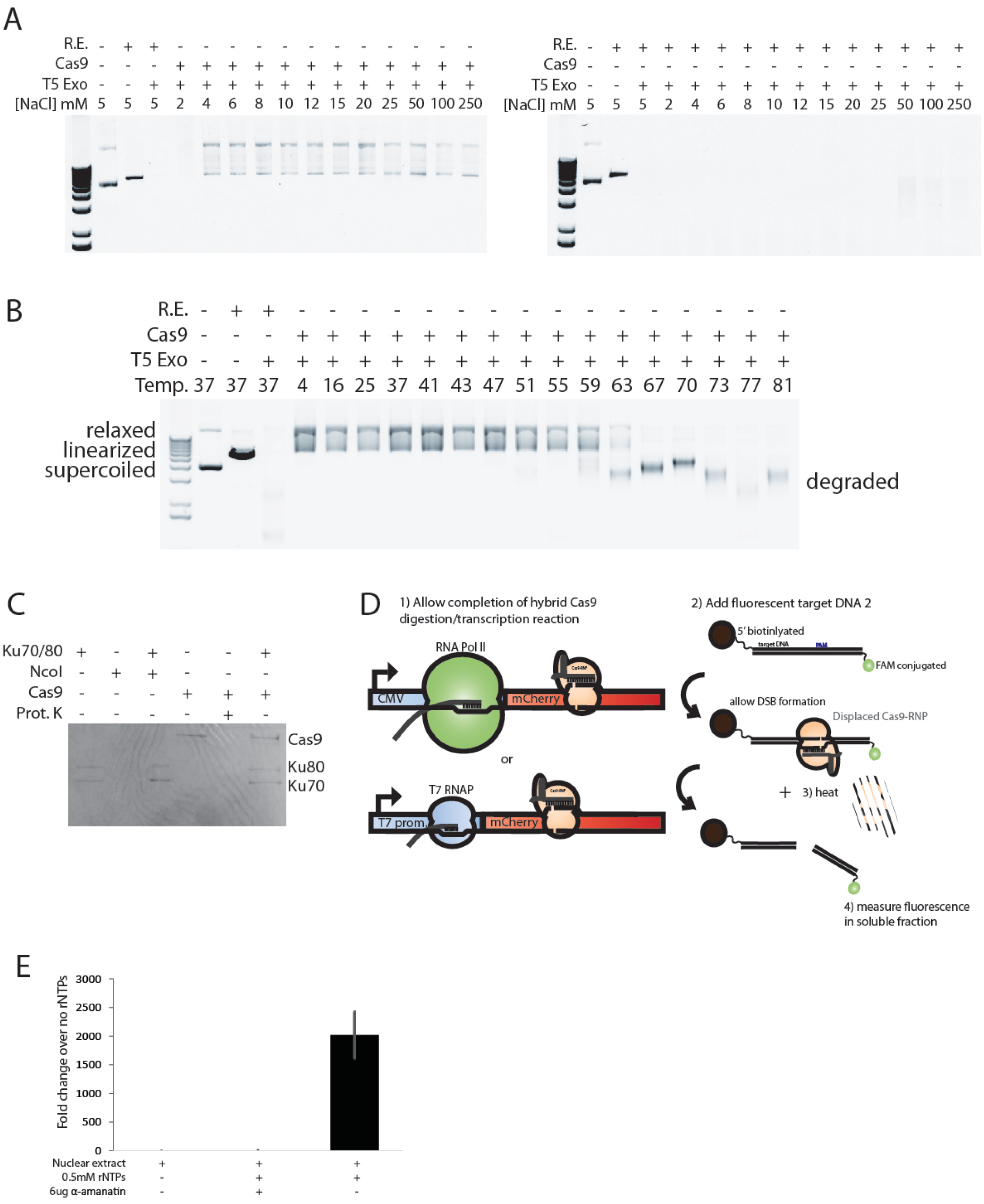
Related to Figure 2 and 3: *In vitro* experiments challenging the Cas9-DSB complex. **A,** (Left) EtBr stained agarose gel showing effects of NaCl on Cas9 protection of DSB. Reactions containing Cas9 were incubated, then subjected to NaCl titrations and mixed. H2O was added to dilute the salt concentration down to 10mM, then T5 exonuclease was added and incubated at 37C for 15m. (Right) Agarose gel showing PmeI restriction endonuclease (R.E) –digested DNA subjected to the aforementioned salt and T5 exonuclease treatments. **B,** Agarose gel showing effect of heat on Cas9 protection of DSB. Cas9-RNP was incubated with target plasmid then shifted to the indicated temperature for 5m. Reactions were cooled to RT then T5 exonuclease was added. Control reactions digested with PmeI (R.E.) demonstrate activity of T5 exonuclease. **C,** Representative Coommassie-stained SDS-PAGE analysis of proteins precipitated with DNA-biotin-streptavidin beads for samples used in Fig 2E. **D,** Schematic illustrating experiment to detect T7 RNAP displaced Cas9 molecules. Target DNA lacks T7 promoter sequence (data shown in Fig 1G). Target DNA 1 and target DNA 2 each have the same DNA sequence targeted by Cas9. Target DNA has a T7 promoter. Target DNA 2 does not have a promoter. **E,** Transcriptional activity of mammalian nuclear extracts. Quantitative qPCR was used to measure mCherry RNA expression in in vitro transcription assays used in Fig 3F. *In vitro* transcription reactions were activate in the presence of 0.5mM rNTPs and inhibited with 6ug/ml α-amanitin.

**Figure S3.**
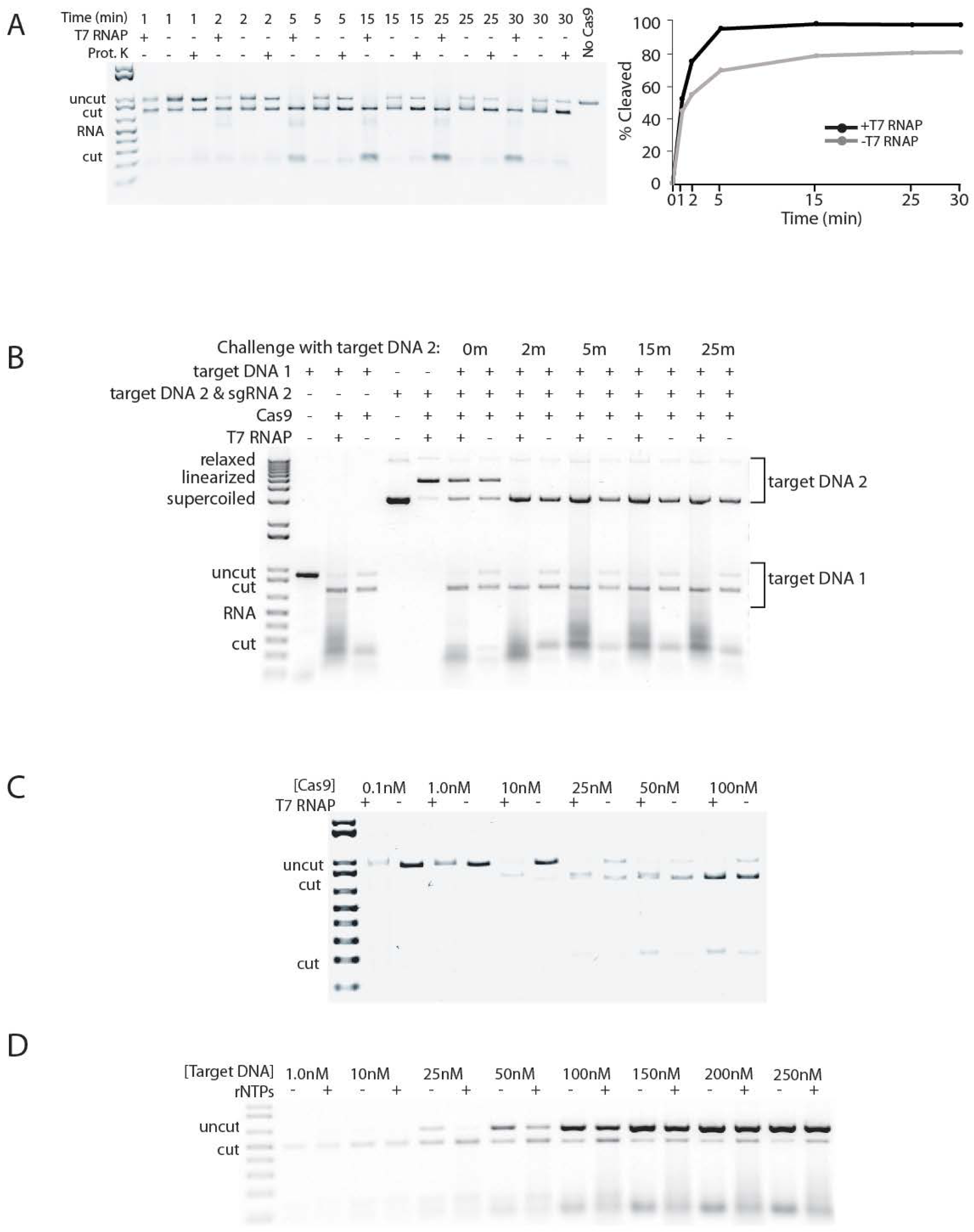
Related to Figure 4: T7 RNAP evicts Cas9 from DSBs, converting it to a multi-turnover nuclease. **A,** Left: Representative agarose gel showing Cas9 digestion reaction in the presence or absence of T7 RNAP over time [Cas9] was 100nM, [Target DNA] was 200nM. Right: quantification of cleavage products over time. **B,** Agarose gel displaying that multi-turnover Cas9 does not release its sgRNA after T7 RNAP mediated removal from DSBs. A linear target DNA (target DNA 1) was incubated with Cas9 and corresponding sgRNA in the presence or absence of T7 RNAP over 25m. A second target DNA lacking the sequences targeted by the first sgRNA (plasmid, target DNA 2) and a new sgRNA targeting DNA 2 were added to the active Cas9 digestion reactions containing target DNA 1 at indicated times. **C,** Representative agarose gel (n = 3) for titration of Cas9 in the presence or absence of T7 RNAP (Fig 4B). Target DNA was held constant at 150nM. **D,** Representative agarose gel (n = 2) for Cas9 digestion reactions with titrations of a target DNA in the presence or absence of T7 RNAP (Fig 4C). Cas9 was held constant at 12.5nM.

**Figure S4.**
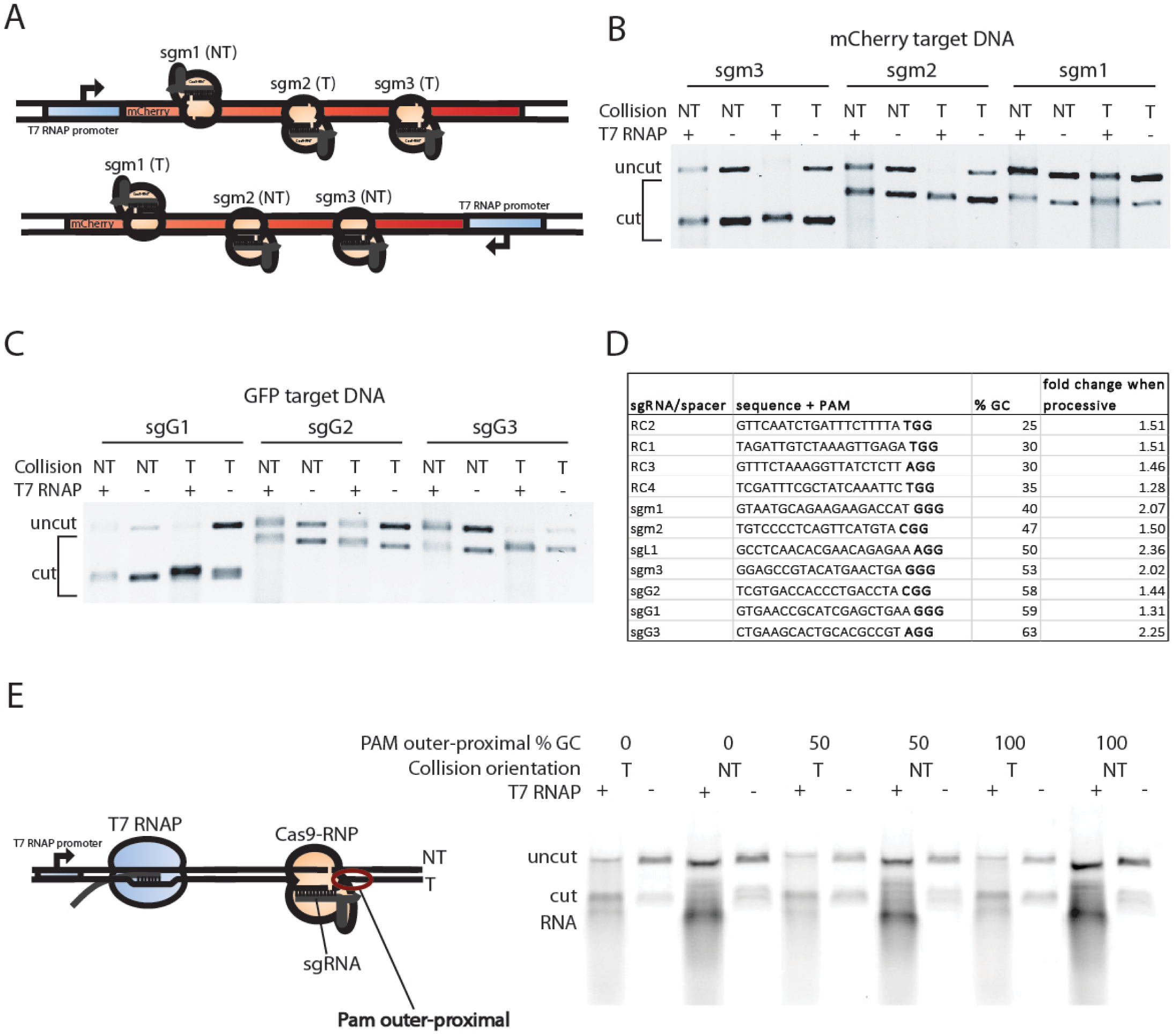
Related to Figure 4: Template strand bias of dislodging Cas9 for multiple sgRNA. **A**, Schematic depicting mCherry target DNA converted into transcription templates with the T7 promoter placed on either end of the DNA to create template or non-template collisions for sgRNA presented in Fig 4D. **B,C**, Representative agarose gel (n = 3) displaying Cas9 digestion reactions presented in Fig 4D. [Cas9] was 100nM, [DNA] was 200nM. **D**, Table containing sgRNA sequences tested for multi-turnover activity shown in Fig 4D and related information. % GC of the sgRNA sequences tested lacks a strong correlation with multi-turnover Cas9 efficiency levels (Pearson correlation: 0.36). **E**, Schematic and agarose gel showing that GC content of the outer-proximal PAM sequence did not effect the lack of T7 RNAP eviction of Cas9 in the non-template orientation. 10bp immediately outer to the PAM were modified to contain varying GC content and used as target DNAs harboring the T7 promoter on either end respectively. Cas9 digestion reactions using sgL1 were performed against these target DNAs in the presence or absence of T7 RNAP. [Cas9] was 100nM and [DNA] was 200nM.

**Figure S5.**
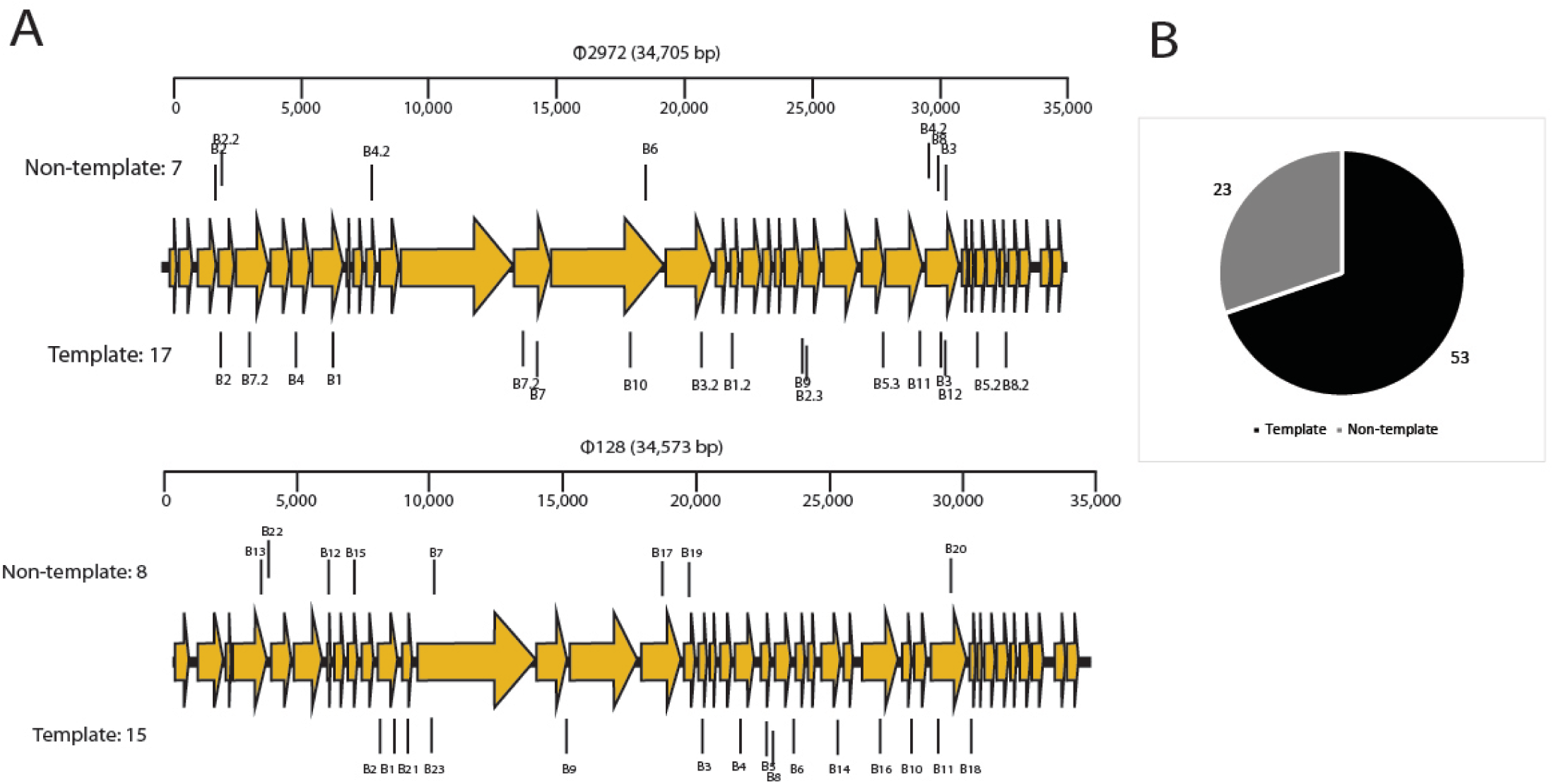
Related to Figure 5: Phage genomes naturally harbor more Cas9 PAM sequences on their template strands. **A,** 68% of Φ128 resistant strains harbor template targeting spacers (Achigar et al, 2017). **B,** 70% of protospacers with unique phage targets map to template strands of various phages. Representative protospacer pool from 15 *S. thermophilus* BIMs (spacer information presented in Table S3).

**Figure S6.**
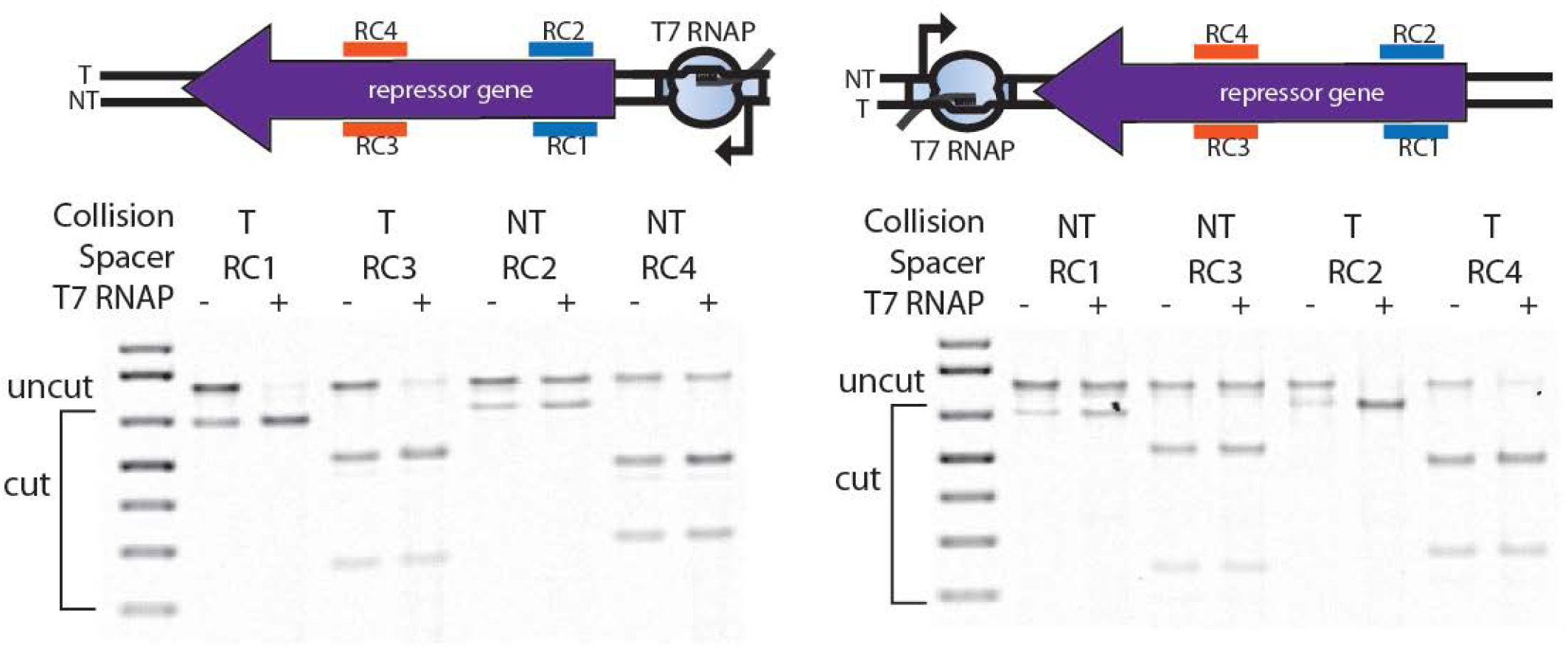
Related to Figure 4 and 6: *In vitro* Cas9 digestion reactions of ΦNM1 with spacers RC1-4 in the presence or absence of T7 RNAP. The ΦNM1 repressor gene was amplified to contain the T7 promoter on either end respectively and subject to Cas9 digestion with spacers RC1-4 in the presence or absence of T7 RNAP. Representative agarose gels display the digestion reactions with and 100nM Cas9 in the presence of 200nM target DNA. Figure 4D incorporates measurements from this experiment, which were performed in duplicate

**Table S1.**
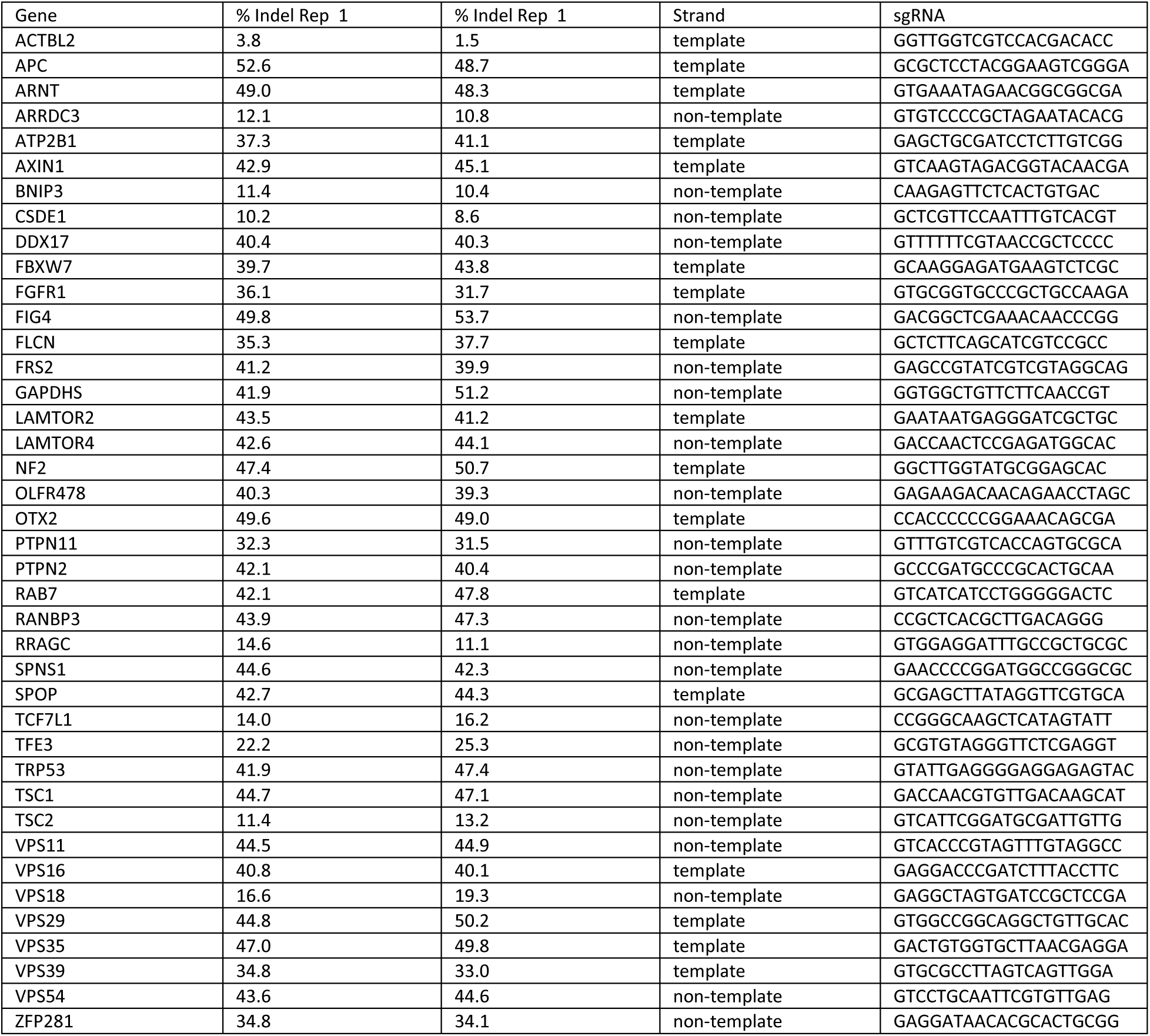
Related to Figure 1: Target gene name, sgRNA sequence, targeted strand, and associated indel frequency for the 40 targeted and deep sequenced mouse genes.

**Table S2.**
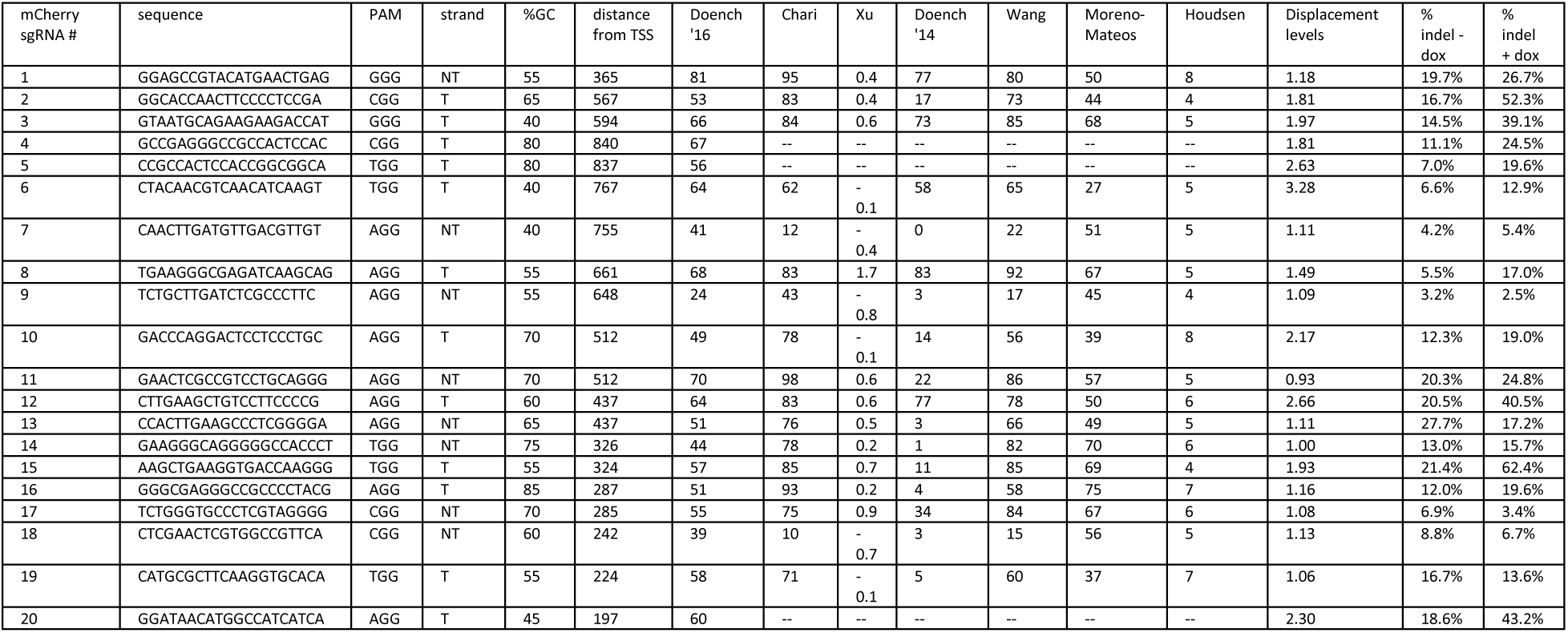
Related to Figure 1: mCherry sgRNA sequences, predicted efficiency scores, indel frequencies and other relevant information to the target sites.

**Table S3.**
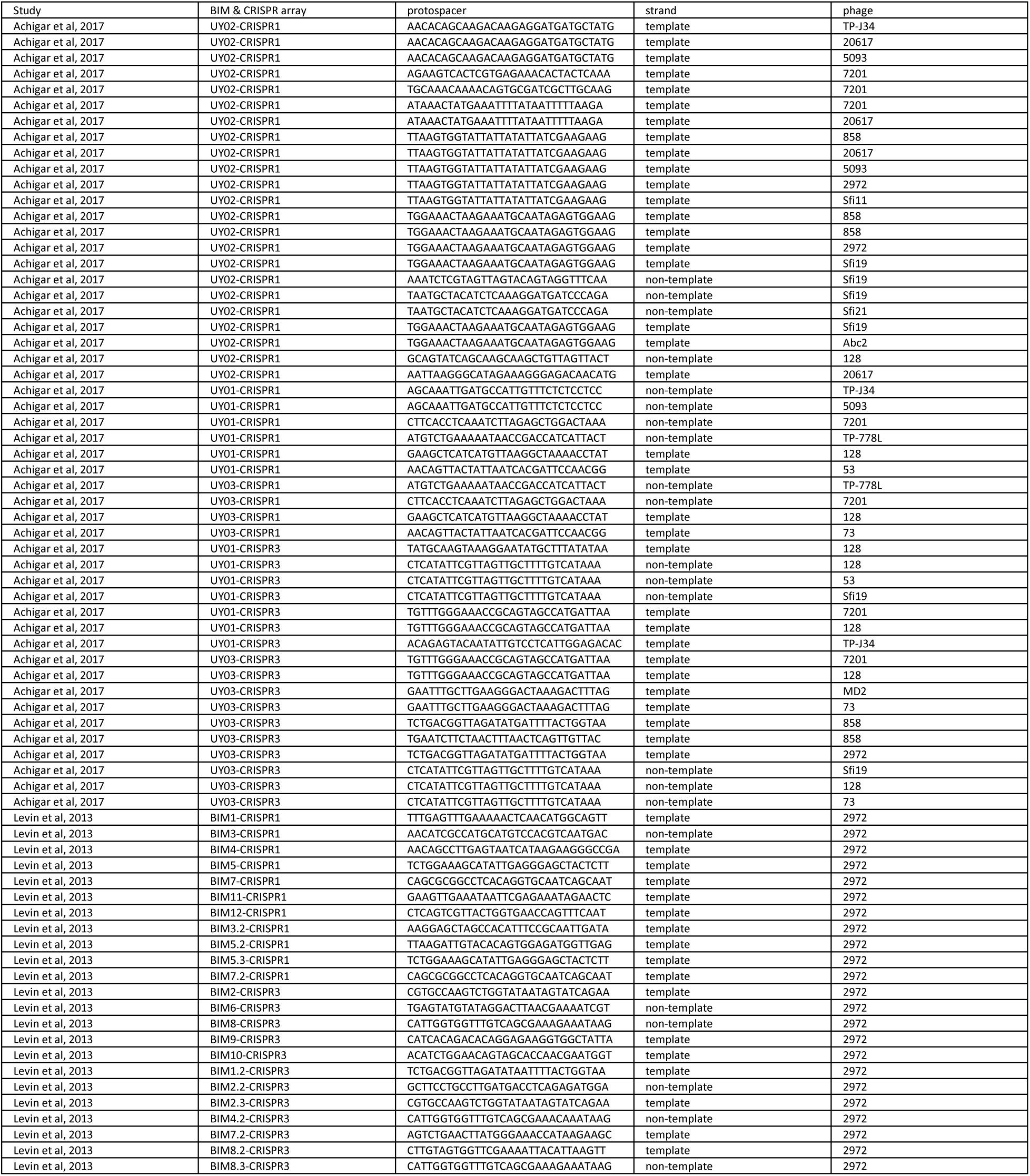
Related to Figure 5, 6, and S5: Bacteriophage insensitive mutant (BIM) protospacer sequences and the strand targeted within each respective phage genome.

